# Pharmacologic Stabilization of Retromer Rescues Endosomal Pathology Induced by Defects in the Alzheimer’s gene *SORL1*

**DOI:** 10.1101/2022.07.31.502217

**Authors:** Swati Mishra, Allison Knupp, Chizuru Kinoshita, C. Andrew Williams, Shannon E. Rose, Refugio Martinez, Panos Theofilas, Jessica E. Young

## Abstract

The Sortilin-related receptor 1 gene (*SORL1,* SORLA) is strongly associated with risk of developing Alzheimer’s disease (AD). SORLA is a regulator of endosomal trafficking in neurons and interacts with retromer, a complex that is a ‘master conductor’ of endosomal trafficking. Pharmacological chaperones stabilize retromer *in vitro*, enhancing its function. Here we used an isogenic series of human induced pluripotent stem cell (hiPSC) lines with either one or two copies of *SORL1* or harboring one copy of a *SORL1* variant linked to increased risk for AD. We treated hiPSC-derived cortical neurons with the established retromer chaperone, TPT-260, and tested whether indicators of AD’s defining endosomal, amyloid, and Tau pathologies were corrected. We observed that the degree of rescue by TPT-260 treatment varied, depending on the number of copies of functional *SORL1* and which *SORL1* variant was expressed. Using a disease-relevant preclinical model, our work illuminates how the *SORL1*-retromer pathway can be therapeutically harnessed.

## Introduction

Alzheimer’s disease (AD) is a devastating neurodegenerative disorder with only a very few medications that alleviate symptoms and no treatment that effectively modifies the course of the disorder for more than several months. Most AD therapies target beta-amyloid (Aβ), the main component of senile plaques, a hallmark AD neuropathology. However, genetic studies, including large genome-wide association studies, have identified dozens of AD risk loci that map to various cellular processes including endo-lysosomal trafficking(Karch and Goate, 2015). Abnormal endosomes and lysosomes in human brains have long been identified as a pathologic hallmark of Alzheimer’s disease(Cataldo et al., 1996; Cataldo et al., 2004; Cataldo et al., 2000). More recent cell biological studies have particularly implicated AD-related deficiencies in trafficking and recycling related to the multi-protein complex retromer, a ‘master conductor’ of endosomal trafficking (Knupp et al., 2020; Simoes et al., 2021; Young et al., 2018). Central to this mechanism is the gene *SORL1*, an endosomal sorting receptor that has emerged as a highly pathogenic AD gene(Scheltens et al., 2021). Missense variants or frameshift variants leading to premature stop codons in *SORL1* can contribute to AD pathogenesis through loss of function of *SORL1(*Pottier et al., 2012; Vardarajan et al., 2014). The *SORL1* gene encodes the sorting receptor SORLA, which engages with retromer as an adaptor protein for multiple cargo, including APP, neurotrophin receptors, and glutamate receptors(Fjorback et al., 2012; Mishra et al., 2022; Simoes et al., 2021). *SORL1* KO neurons derived from hiPSCs have endosomal traffic jams, mislocalized neuronal cargo and show impairments in endosomal degradation and recycling(Hung et al., 2021; Knupp et al., 2020; Mishra et al., 2022). Both endosomal recycling and degradation pathways in neurons are enhanced in a model where *SORL1* is overexpressed(Mishra et al., 2022), suggesting that increasing *SORL1* expression in neurons may be beneficial in AD. However, *SORL1* is a difficult therapeutic target due to its size: the gene contains 48 exons and large intronic regions and the protein itself is 2214 amino acids(Rogaeva et al., 2007).

Enhancing endosomal trafficking via targeting the retromer complex, which is intimately associated with *SORL1*, may be a feasible therapeutic strategy. Small molecule pharmacologic chaperones that stabilize the cargo recognition core of retromer have been developed and reduce amyloidogenic processing of APP in a mouse model of AD and increase the flow of SORLA through endosomes(Mecozzi et al., 2014). Retromer chaperones have been shown to reduce Aβ and phospho-Tau in hiPSC-cortical neurons from AD and controls and promote neuroprotection in hiPSC-motor neurons from amyotrophic lateral sclerosis (ALS) patients (Muzio et al., 2020; Young et al., 2018). However, how they affect other defining pathologies of AD (i.e.: endosomal pathologies) has not been shown.

We used our previously published *SORL1* deficient hiPSC cell lines (*SORL1* KO)(Knupp et al., 2020) and new lines engineered to have loss of *SORL1* on one allele (*SORL1*^+/-^) or to have one copy of an AD-associated *SORL1* variant (*SORL1*^Var^) to investigate whether enhancing retromer-related trafficking using TPT-260, an established retromer chaperone(Chu and Pratico, 2017; Mecozzi et al., 2014; Vagnozzi et al., 2021), can improve endosomal phenotypes. We report that in neurons derived from *SORL1*^+/-^ and *SORL1*^Var^ lines, we observe enlarged early endosomes, altered endosomal localization of cellular cargo and deficits in endosomal recycling. We show that treatment with TPT-260 rescues these phenotypes, although the degree of rescue of these endosomal phenotypes is variable, depending on which *SORL1* variant was tested and the number of functional copies of *SORL1* were present. Recent work has begun to suggest that *SORL1* variants may differ in their pathogenicity(Holstege et al., 2022) and that different classes of *SORL1* variants may emerge(Andersen, 2023). Our data supports the idea that there may be different mechanisms of pathogenicity for SORL1 variants.

We also show that in addition to rescuing endo-lysosomal phenotypes, TPT-260 lowers Aβ and phosphorylated Tau (p-Tau) in all *SORL1* conditions, suggesting that enhancing retromer can improve multiple cellular phenotypes in AD. Loss of one copy of *SORL1* has been shown to be causative for AD(Andersen et al., 2022; Scheltens et al., 2021) and, in light of pathogenic variants in *SORL1* that continue to be identified(Fazeli et al., 2023; Jensen AM, 2023), our data suggest that the *SORL1*-retromer axis is important for future therapeutic development in AD.

## Results

### *SORL1*^+/-^ and *SORL1*^Var^ hiPSC-derived neurons have swollen early endosomes and increased Aβ secretion

Variants in the VPS10 domain of *SORL1* have been shown to be damaging(Holstege et al., 2017) and have been identified in early-onset AD families that do not have familial AD mutations in the amyloid precursor protein (*APP*) genes or in the presenilin 1/presenilin 2 genes (*PSEN1/2*)(Pottier et al., 2012). The VPS10 domain of the SORL1 protein, SORLA, is characterized as a ligand binding domain and has been shown to bind amyloid beta (Aβ) peptides(Caglayan et al., 2014). Using CRISPR/Cas9 genome editing, we generated isogenic human induced pluripotent stem cell (hiPSC) lines containing either heterozygous AD-associated variants in the VPS10 domain: E270K, Y141C, and G511R *(SORL1*^Var^) or heterozygous knock-out *SORL1* (*SORL1*^+/-^) **(Figure S1).** These variants have all been associated with increased AD risk(Caglayan et al., 2014; Pottier et al., 2012; Vardarajan et al., 2014), although the E270K variant has been found in control cases(Holstege et al., 2022). Western blot analysis revealed that *SORL1*^Var^ neurons do not have a loss of SORLA protein expression as these missense variants do not introduce a frame-shift or destabilize the protein enough for it to be degraded **(Figure S1**). *SORL1*^+/-^ neurons have 50% of SORLA expression compared with isogenic WT controls (**Figure S1).**

We and others have previously observed enlarged early endosomes in homozygous *SORL1* KO hiPSC-derived neurons(Hung et al., 2021; Knupp et al., 2020). We hypothesized that *SORL1*^+/-^ and *SORL1*^Var^ neurons would similarly contain enlarged early endosomes, although we predicted that due to the single copy of WT *SORL1*, the phenotype could be more subtle. We immunostained *SORL1* KO, *SORL1*^+/-^, and *SORL1*^Var^ neurons for EEA1 to mark early endosomes, imaged cells using confocal microscopy, and quantified early endosome size as we have previously described(Knupp et al., 2020). We observed a significant increase in endosome size in *SORL1*^Var^, *SORL1*^+/-^, and *SORL1* KO neurons compared to isogenic WT controls (**Figure 1 A-B**). Interestingly, endosomes in neurons with one copy of WT *SORL1* (*SORL1*^Var^ and *SORL1*^+/-^) were not as enlarged as in cells fully deficient in *SORL1* (*SORL1* KO) (**Figure 1B**).

**Figure 1.**
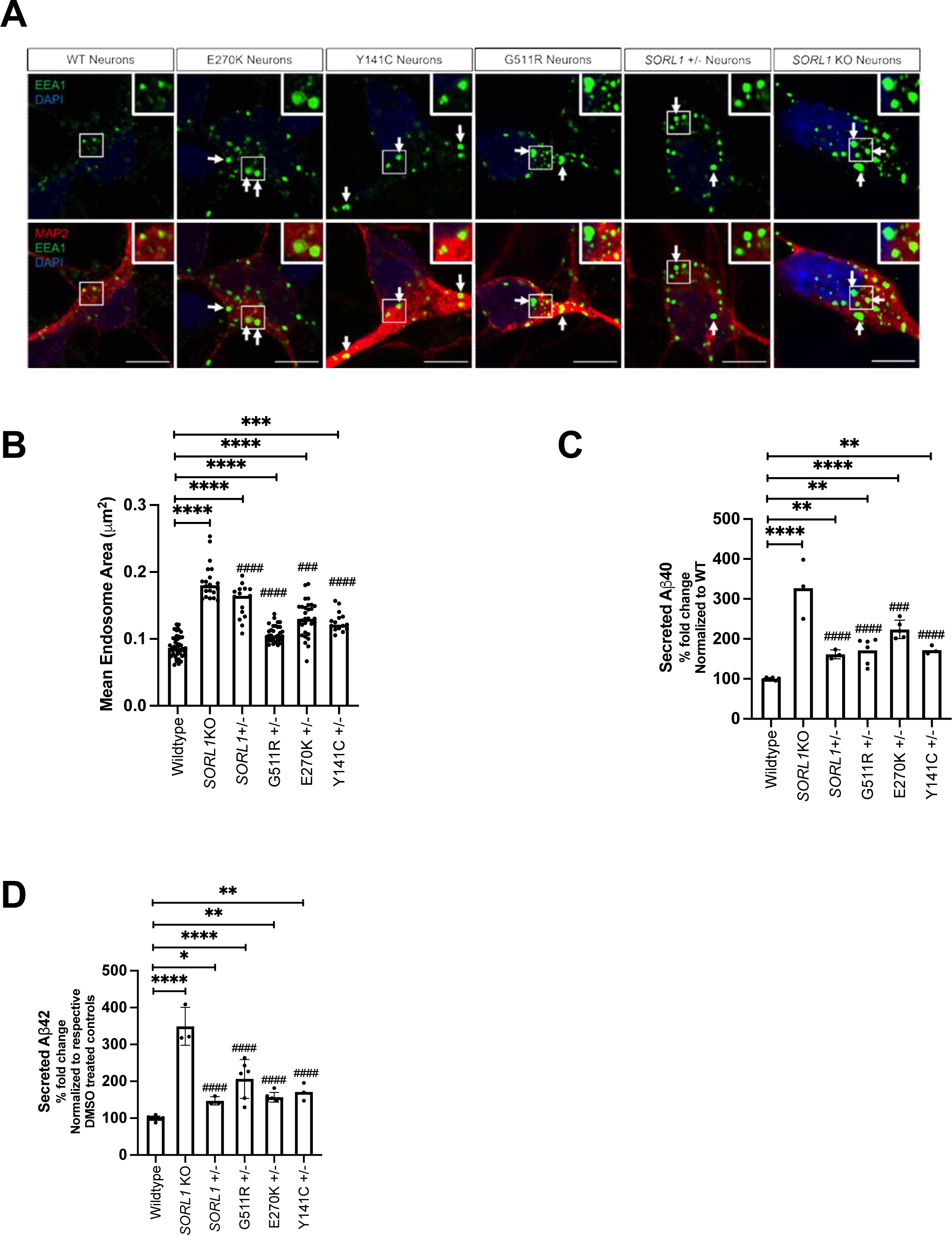
*SORL1^+/-^*and *SORL1^Var^* hiPSC-derived neurons have swollen early endosomes and increased Aβ secretion. **(A)** Representative immunofluorescent images of Wildtype (WT), heterozygous *SORL1_Var_*, *SORL1*_+/-_, and *SORL1* KO hiPSC-derived neurons. Scale bar: 5 μm. **(B)** The size of early endosome marker, EEA1 puncta is larger in *SORL1*_Var_, *SORL1*_+/-_, and *SORL1* KO neurons than in WT neurons, indicated by asterisks. Each datapoint on the graph represents mean early endosome area/image. The difference between *SORL1^+/-^* and *SORL1^Var^* as compared to *SORL1* KO is indicated by hashmarks. **(C-D) Heterozygous** *SORL1*^Var^, *SORL1*^+/-,^ and *SORL1* KO hiPSC-derived neurons secrete increased levels of **(C)**Aβ_40_ and **(D)**Aβ_42_ as compared to WT controls (asterisks). *SORL1* KO neurons secrete increased levels of Aβ_40_ and Aβ_42_ as compared to *SORL1*_Var_ and *SORL1*_+/-_ hiPSC-derived neurons. (hashmarks). For imaging experiments, 10-15 images and 1-3 clones were analyzed per genotype. For Aβ secretion experiments, 1-3 clones and 3 replicates per clone per genotype were used. Each data point represents Aβ levels measured from the media per clone/per well. Data represented as mean ± SD. Data was analyzed using parametric one-way ANOVA. Significance was defined as a value of */^#^p < 0.05, **/^##^p < 0.01, ***/^###^p < 0.001, and ****/^####^p < 0.0001. ns= not significant

When SORLA is absent, APP is unable to be trafficked from early and recycling endosomes where it is more readily processed to Aβ(Das et al., 2016; Knupp et al., 2020; Mishra et al., 2022; Tan and Gleeson, 2019; Toh et al., 2018). We have previously reported increased secreted Aβ in *SORL1* KO neurons(Knupp et al., 2020) and here we observed an increase in secreted Aβ peptides in *SORL1*^+/-^ or *SORL1*^Var^ neurons compared to WT neurons. Consistent with having one functional copy of *SORL1*, the increase in Aβ was not as pronounced as with the full *SORL1* KO (**Figure 1C-D**).

### TPT-260 treatment increases VPS26B expression and reduces endosome size in *SORL1* **KO, *SORL1*^+/-^, and *SORL1*^Var^ neurons.**

We next tested whether enhancing endo-lysosomal function via small molecules could rescue pathological phenotypes. We chose to study the effects of a pharmacological chaperone, TPT-260 (also called R55), which stabilizes and enhances retromer by binding the VPS35-VPS26-VPS29 trimer in the cargo-recognition core of the multi-subunit retromer complex(Mecozzi et al., 2014). Previously, we have shown that a similar molecule, TPT-172 (also called R33), was effective at reducing Aβ and pTau levels in hiPSC-derived neurons from AD patients and controls(Young et al., 2018). First, we tested whether TPT-260 treatment enhances expression of retromer components. SORLA directly interacts with VPS26(Fjorback et al., 2012), therefore we analyzed VPS26B protein expression as this component of the retromer is an isoform that is enriched in neurons(Simoes et al., 2021). We show that in all conditions (*SORL1* WT, *SORL1* KO, *SORL1*^+/-^, and *SORL1*^Var^) treatment with TPT-260 significantly increases VPS26B protein expression (**Figure 2A-B**). We next examined whether TPT-260 treatment influences early endosome size. In *SORL1* KO, *SORL1*^+/-^, and *SORL1*^Var^ neurons, treatment with TPT-260 reduced endosome size compared with vehicle (DMSO) treated controls (**Figure 2C-D**). In TPT-260 treated *SORL1^+/-^*and *SORL1^Var^* neurons, endosome size was not significantly different than in WT neurons treated with TPT-260. However, while endosome size was also reduced in neurons with full *SORL1* KO, it was not reduced to WT levels, indicating that without at least one copy of *SORL1*, this phenotype cannot be fully resolved (**Figure 2D**). We did not observe any changes in EEA1 puncta size in WT neurons treated with TPT260, indicating that the retromer chaperone does not alter the size of endosomes that are not enlarged.

**Figure 2.**
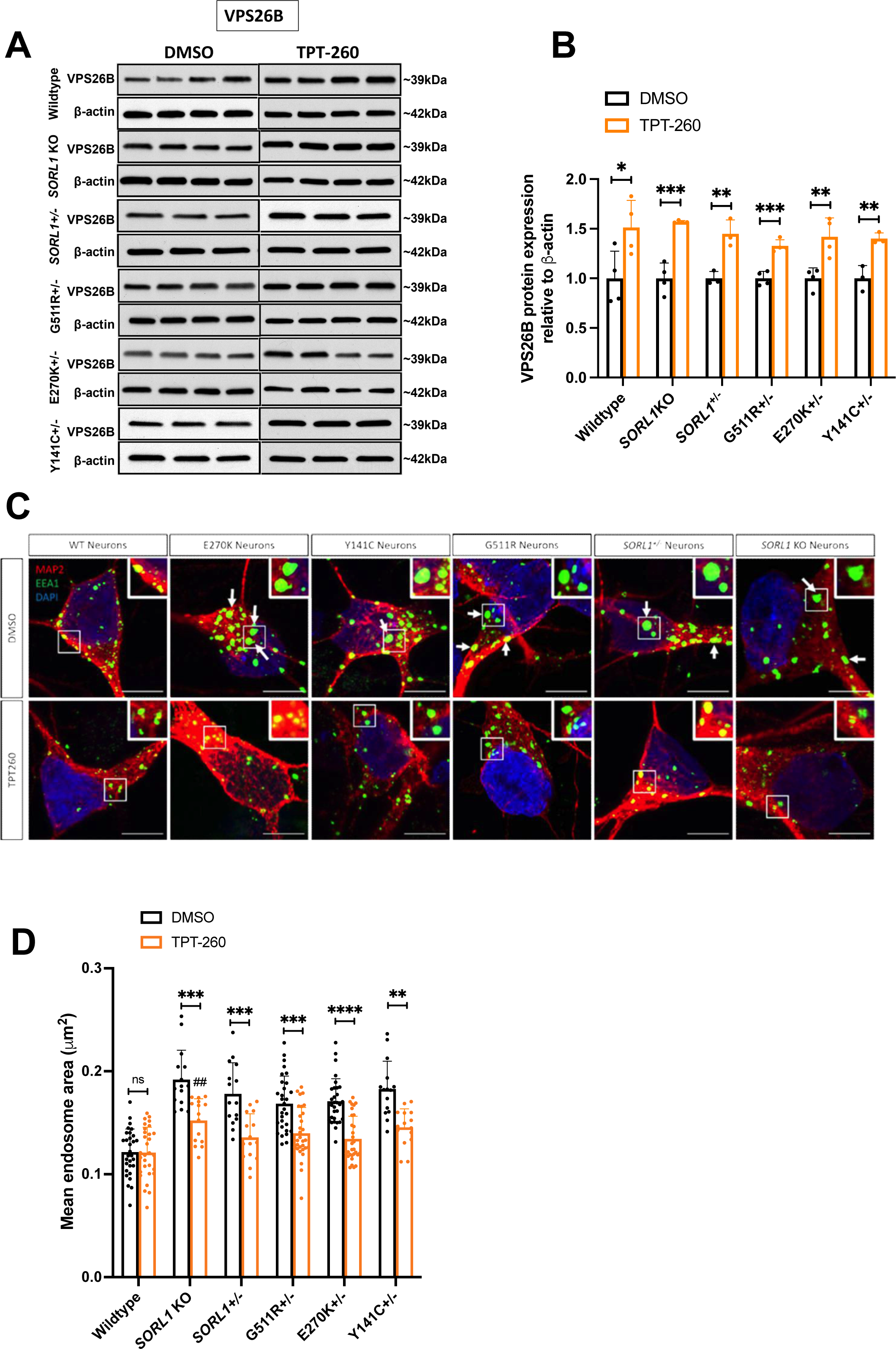
Retromer enhancement with TPT-260 rescues enlarged early endosome phenotype. **(A-B)** Western blot showing increased protein expression of retromer subunit VPS26B upon TPT-260 treatment of *SORL1_Var_*, *SORL1_+/-_* and *SORL1* KO hiPSC-derived neurons. **(A)** Representative western blot images of DMSO and TPT-260 treated hiPSC-derived neurons. **(B)** Quantification of protein expression of VPS26B observed in (A) using Image J. For this experiment, N=4; 2 clones/genotype of WT, *SORL1* KO, G511R+/- and E270K+/- and 2 replicates/clone were used. For Y141C+/- and SORL1+/- cell lines, 1 clone/genotype and 3 replicates/clone were used. **(C)** Representative immunofluorescent images of TPT-260 and DMSO-treated WT, heterozygous *SORL1_Var_*, *SORL1_+/-_* and *SORL1* KO hiPSC-derived neurons. Scale bar: 5 μm. **(D)** TPT-260 treatment reduces endosome size in *SORL1*^Var^, *SORL1*^+/-^, and *SORL1* KO neurons (indicated by asterisks). Endosome size in TPT-260 treated *SORL1* KO neurons is significantly different from *SORL1*^Var^ and *SORL1*^+/-^ neurons (indicated by hashmarks). Each datapoint on the graph represents mean early endosome area/image. N=10-20 images analyzed per genotype. Data represented as mean ± SD. For all experiments, 1-3 clones and 3 replicates per clone per genotype were used. Normally distributed data was analyzed using parametric two-way ANOVA. 1-3 clones were analyzed per genotype. Significance was defined as a value of */^#^p < 0.05, **/^##^p < 0.01, ***/^###^p < 0.001, and ****/^####^p < 0.0001. ns= not significant

### Retromer enhancement reduces secreted Aβ levels and pTau levels in *SORL1* KO, *SORL1*^+/^, and *SORL1*^Var^ neurons

We also determined that TPT-260 treatment reduces Aβ peptides in *SORL1* KO, *SORL1*^+/-^ or *SORL1*^Var^ neurons. From TPT-treated cultures we measured secretion of Aβ peptides into culture medium. In all cell lines we observed decreased secreted Aβ peptides. In TPT-260 treated neurons with full *SORL1* KO, Aβ 1-40 levels were significantly higher than in the other treated conditions, although still reduced from non-treated conditions. (**Figure 2A**). Interestingly, Aβ 1-42 levels were not changed in *SORL1* KO neurons although they were reduced to WT levels in all cell lines with one copy of WT *SORL1* (**Figure 2B**).

Accumulation of phosphorylated Tau (p-Tau) protein is a significant neuropathological hallmark in AD and retromer chaperones have been reported to decrease p-Tau levels in both hiPSC-neurons and mouse models of AD and tauopathy(Li et al., 2020; Young et al., 2018). In our hiPSC-neuronal model, we did not detect significant changes in p-Tau in neurons without *SORL1* or in any of our *SORL1*^Var^ neurons (**Figure S2 A-C**), although in other studies where *SORL1* KO hiPSC-derived neurons are differentiated using the expression of the transcription factor NGN2, loss of *SORL1* does lead to increased p-Tau(Lee et al., 2023). Despite this, when we treated all genotypes of neurons with TPT-260, we observed that treatment did significantly reduce p-Tau at several epitopes and there was no difference in the level of reduction relative to whether neurons were fully deficient in *SORL1* or harbored an AD risk variant (**Figure 3C-E**).

**Figure 3.**
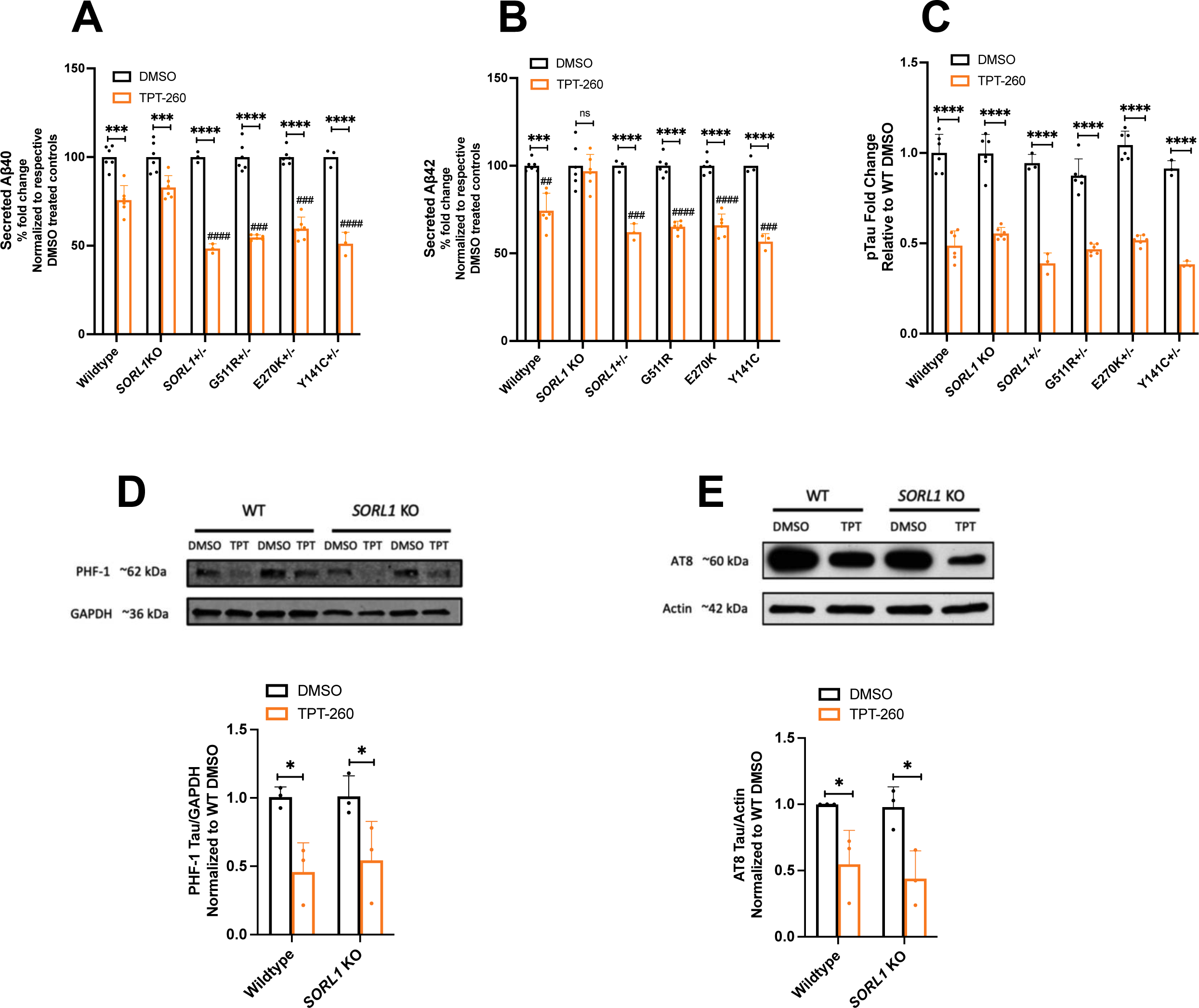
Retromer enhancement with TPT-260 reduces AD pathological phenotypes. **(A-B)** Levels of secreted Aβ 1-40 **(A)** and Aβ 1-42 **(B)** are reduced with TPT-260 treatment in all genotypes, compared to DMSO, except secreted Aβ 1-42 in *SORL1* KO neurons (indicated by asterisks). The % decrease in Aβ 1-40 and Aβ 1-42 levels of TPT-260 treated cells is greater in *SORL1^Var^* and *SORL1*^+/-^ neurons as compared to *SORL1* KO neurons (indicated by hashmarks). For all experiments, 1-3 clones and 3 replicates per clone per genotype were used. Each data point represents Aβ levels measured from the media per clone/per well. Data represented as mean ±SD. (**C-E**) Tau phosphorylation on three epitopes was examined in response to TPT-260 treatment. **(C)** Thr 231 epitope levels, as measured by ELISA assay, is reduced in all the genotypes treated with TPT-260 relative to WT DMSO controls. Each datapoint represents total tau or phospho tau levels reassured from the cell lysate per clone/per well. **(D)** The PHF-1 (Ser396/Ser404) and the **(E)** AT8(Ser202/Thr305) epitopes as measured by western blot is decreased in TPT-260 treated *SORL1* KO hiPSC-derived neurons relative to WT DMSO controls. 1-3 clones were analyzed per genotype. Each datapoint represents total tau or phospho tau levels reassured from the cell lysate per clone/per well. Data represented as mean ± SD. Normally distributed data was analyzed using parametric two-way ANOVA. 1-3 clones were analyzed per genotype. Significance was defined as a value of */^#^p < 0.05, **/^##^p < 0.01, ***/^###^p < 0.001, and ****/^####^p < 0.0001. ns= not significant

### Lysosomal stress and degradation are improved by pharmacologic stabilization of retromer

We previously demonstrated that SORL1 KO neurons also have enlarged lysosomes and impaired lysosomal degradation(Mishra et al., 2022). Here we examined lysosome size in SORL1^+/-^ and SORL1^Var^ neurons. Interestingly, in SORL1^+/-^ and in two of the variant lines (Y141C and E270K) we did not observe enlarged lysosomes (**Figure 4 A-B**). However, in a different variant cell line, G511R, we did document a significantly increased lysosome size that is reduced after TPT-260 treatment (**Figure 4C-D**). To test whether impaired lysosomal degradation in SORL1 KO neurons can be rescued by retromer enhancement, we used the DQ-Red-BSA assay(Marwaha and Sharma, 2017). We observed a significant reduction in DQ-Red-BSA fluorescence, indicating impaired degradation, at 24 hours in DMSO treated *SORL1* KO neurons, consistent with our previous observations(Mishra et al., 2022). In TPT-260 treated neurons, we document a complete rescue of this phenotype (**Figure 4D-E**) suggesting that retromer stabilization can promote this pathway, even in the absence of *SORL1*.

**Figure 4.**
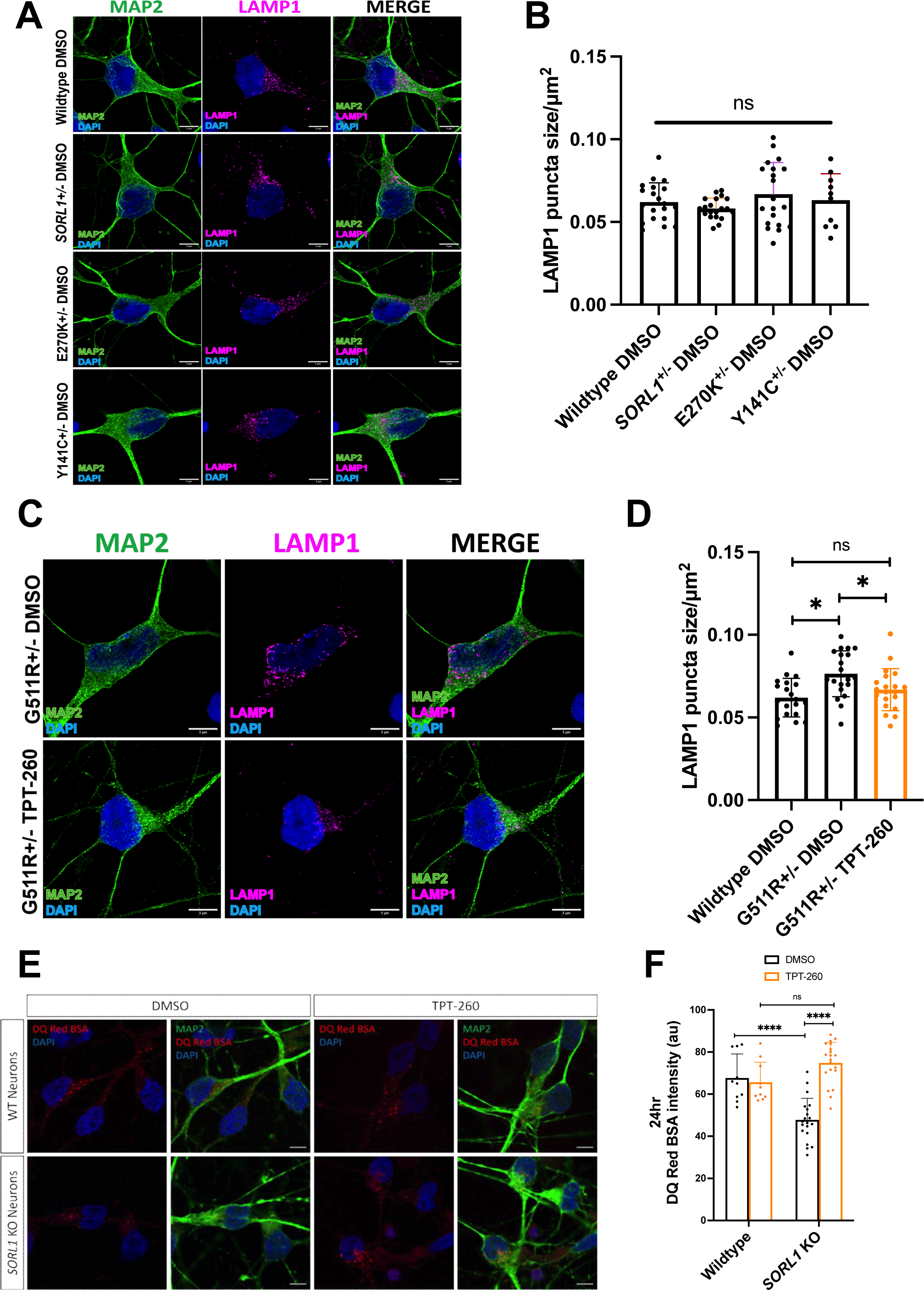
Retromer enhancement with TPT-260 rescues enlarged endosome phenotype in G511R+/- hiPSC-derived neurons and enhances lysosomal degradation in *SORL1* KO neurons. **(A-B)** Lysosome size measured using lysosome specific marker LAMP1 in WT, *SORL1_Var_* and *SORL1_+/-_* hiPSC-derived neurons. **(A)** Representative images showing no difference in lysosome size of *SORL1*+/-, E270K+/- and Y141C+/- hiPSC-derived neurons as compared to WT hiPSC-derived neurons. **(B)** Quantification of lysosome size observed in (A) as indicated by LAMP1 puncta size/μm^2^ using Cell Profiler software. **(C-D)** Retromer enhancement with TPT-260 rescues enlarged lysosome size in G511R+/- hiPSC-derived neuron**s (C)** Representative images showing rescue of enlarged lysosome size in G511R+/- hiPSC-derived neurons and **(D)** Quantification of lysosome size observed in (C) using Cell Profiler software. Each data point on the graphs indicates mean LAMP1 puncta size/image normalized to mean cell area/image. 2 clones/genotype were used for the experiments mentioned above. Scale bar: 5μm **(E-F)** TPT-260 rescues degradation of DQ-Red BSA in *SORL1* KO hiPSC-derived neurons. **(E)** Representative images of WT and *SORL1* KO neurons treated with DQ-Red-BSA for 24 hours. **(F)**TPT-260 rescues lysosomal degradation shown by increased fluorescence intensity of DQ-Red BSA after 24 hrs in TPT-260 treated *SORL1* KO neurons as compared DMSO. There is no difference between DMSO treated and TPT-260 treated WT hiPSC-derived neurons. Scale bar: 5 μm. 1-2 clones/genotype and at least 10 images/clone were used for this experiment. Each datapoint on the graph indicates mean fluorescence intensity of DQ-BSA/image. Data represented as mean ± SD. Normally distributed data was analyzed using parametric two-way ANOVA. 1-3 clones were analyzed per genotype. Significance was defined as a value of */^#^p < 0.05, **/^##^p < 0.01, ***/^###^p < 0.001, and ****/^####^p < 0.0001. ns= not significant

### Endosomal mis-localization of retromer and APP proteins is fully or partially rescued by TPT-260 treatment, depending on whether one WT copy of *SORL1* is present

In *SORL1* KO neurons, both VPS35, a core component of retromer, and APP have increased localization in early endosomes(Knupp et al., 2020; Mishra et al., 2022). We examined whether TPT-260 treatment could correct this mis-localization. By performing colocalization analysis for EEA1 (early endosomes) and VPS35, we observed that *SORL1*^+/-^ neurons also had increased VPS35 accumulation in early endosomes (**Figure 5A-B**). Treatment with TPT-260 completely rescued VPS35/EEA1 colocalization in *SORL1*^+/-^ and *SORL1* KO neurons, indicating that this treatment was able to mobilize retromer away from early endosomes even in the absence of *SORL1* (**Figure 5A-B**). *SORL1*^+/-^ neurons also have increased colocalization of APP with EEA1, however upon TPT-260 treatment APP/EEA1 colocalization was completely resolved in *SORL1*^+/-^ neurons, while there was still significantly more APP in early endosomes in treated *SORL1* KO neurons, even though total APP expression is not different. (**Figure 5C-D, Figure S2D-E**). This observation is consistent with *SORL1* being a main adaptor protein for APP and retromer via VPS26(Fjorback et al., 2012) and could also explain why the reduction of Aβ peptides in TPT-260 treated *SORL1* KO neurons is not as significant as in neurons with at least one WT copy of *SORL1*; in neurons without any copies of *SORL1*, APP is still largely localized to early endosomes where it can be processed to Aβ. Because we observed that treatment with TPT-260 can improve lysosomal degradation in SORL1 KO neurons using DQ-Red-BSA, we tested whether the small reduction in APP co-localization with EEA1 observed in SORL1 KO neurons after TPT-260 treatment could be due to changes in the degradative capacity of lysosomes. We treated *SORL1* KO+TPT-260 neurons with Bafilomycin-A, which disrupts lysosomal acidity, and tested whether the partial rescue of APP co-localization observed with TPT-260 was reversed. We determined that treatment with Bafilomycin-A did indeed impair lysosomal degradation (**Figure S3A**), however it did not affect the co-localization of APP in *SORL1* KO cells(**Figure S3B-C**). This suggests that in *SORL1* KO+TPT-260 neurons, the small reduction of APP in early endosomes is not due to increased lysosomal degradation.

**Figure 5.**
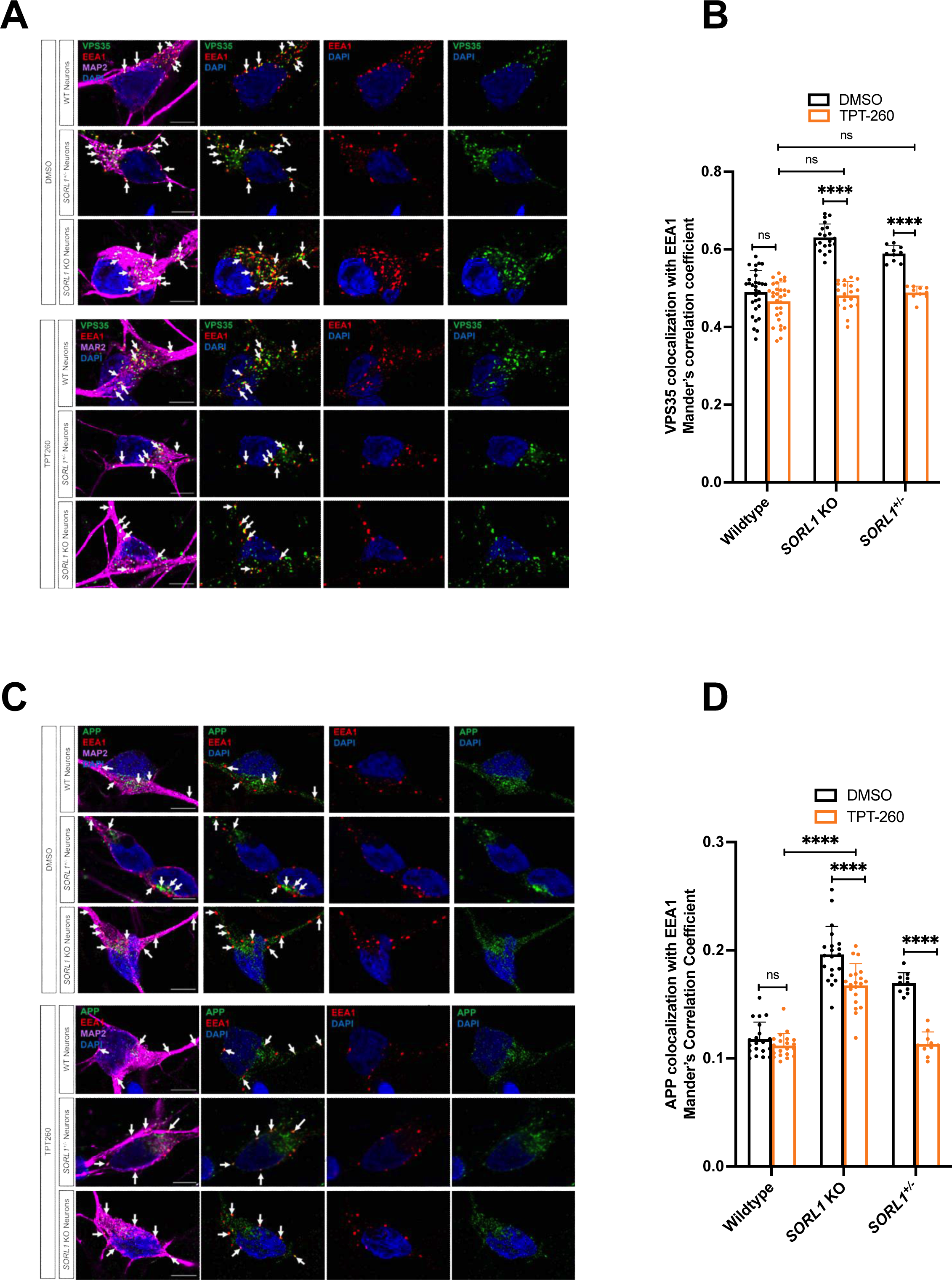
TPT-260 treatment reduces localization of APP and VPS35 in early endosomes. **(A-B)** Co-localization of VPS35 (green) with EEA1 (red) is increased in *SORL1*_+/-_ and *SORL1* KO as compared to WT hiPSC-derived neurons. In *SORL1*_+/-_ and *SORL1* KO neurons, treatment with TPT-260 colocalization of VPS35 and EEA1 is not different than in WT cells (white arrows). **(C-D)** Co-localization of APP (green) with EEA1 (red) co-localization is increased in *SORL1*_+/-_ and *SORL1* KO neurons. Treatment with TPT-260 reduces APP/EEA1 colocalization to a greater extent in *SORL1* _+/-_ neurons such that it is not different from WT neurons treated with TPT-260 (white arrows). However, in *SORL1* KO neurons, APP/EEA1 colocalization is still significantly increased compared to WT cells. Scale bar: 5 μm. For co-localization analysis, 10-15 images per clones and 1-3 clones were analyzed per genotype. Each data point on the graph indicates Mander’s correlation coefficient measured/image using Image J. Data represented as mean ± SD. Data was analyzed using parametric one-way ANOVA. Significance was defined as a value of */^#^p < 0.05, **/^##^p < 0.01, ***/^###^p < 0.001, and ****/^####^p < 0.0001. ns= not significant

### Impaired endosomal recycling in SORL1^+/-^ neurons is improved by pharmacologic stabilization of retromer

In *SORL1* KO neurons, endosomal traffic jams impede cargo traffic to late endosomes and lysosomes as well as to the cell surface recycling pathway(Mishra et al., 2022). The latter function has been shown to involve a neuron-specific subunit of retromer, VPS26B(Simoes et al., 2021). To test endosomal recycling, we utilized the transferrin recycling assay, a measure of both fast and slow endosomal recycling(Ouellette and Carabeo, 2010; Sonnichsen et al., 2000). We found that in *SORL1* KO neurons, TPT-260 treatment only partially rescued endosomal recycling (**Figure 6 A-B**). However, when TPT-260 treated *SORL1^+/-^* neurons were analyzed, we document a complete rescue of transferrin recycling (**Figure 6 A,C**). These data suggest that enhancing retromer can promote endosomal recycling but at least one functional copy of *SORL1* is needed to recycle cargo as efficiently as in WT neurons. We next tested cell surface levels of GLUA1, a subunit of excitatory AMPA receptors and an important protein in neurotransmission. Previous work has demonstrated that in conditions of both *SORL1* and *VPS26B* deficiency, there is a reduction in cell surface levels of GLUA1 subunits in neurons that adversely affects their physiology(Mishra et al., 2022; Simoes et al., 2021). In *SORL1 KO, SORL1*^+/-^ and G511R variant neurons, we observed significantly reduced levels of GLUA1 on the neuronal surface (**Figure 6 D, E**). We did not observe changes in surface GLUA1 levels in Y141C and E270K variant neurons. Treatment with TPT-260 increased levels of GLUA1 on the cell surface in *SORL1* +/- neurons but not in *SORL1* KO neurons (**Figure 6 D, E**). Interestingly, TPT-260 did not rescue cell surface levels of GLUA1 in G511R variant neurons, even with one copy of the WT allele.

**Figure 6.**
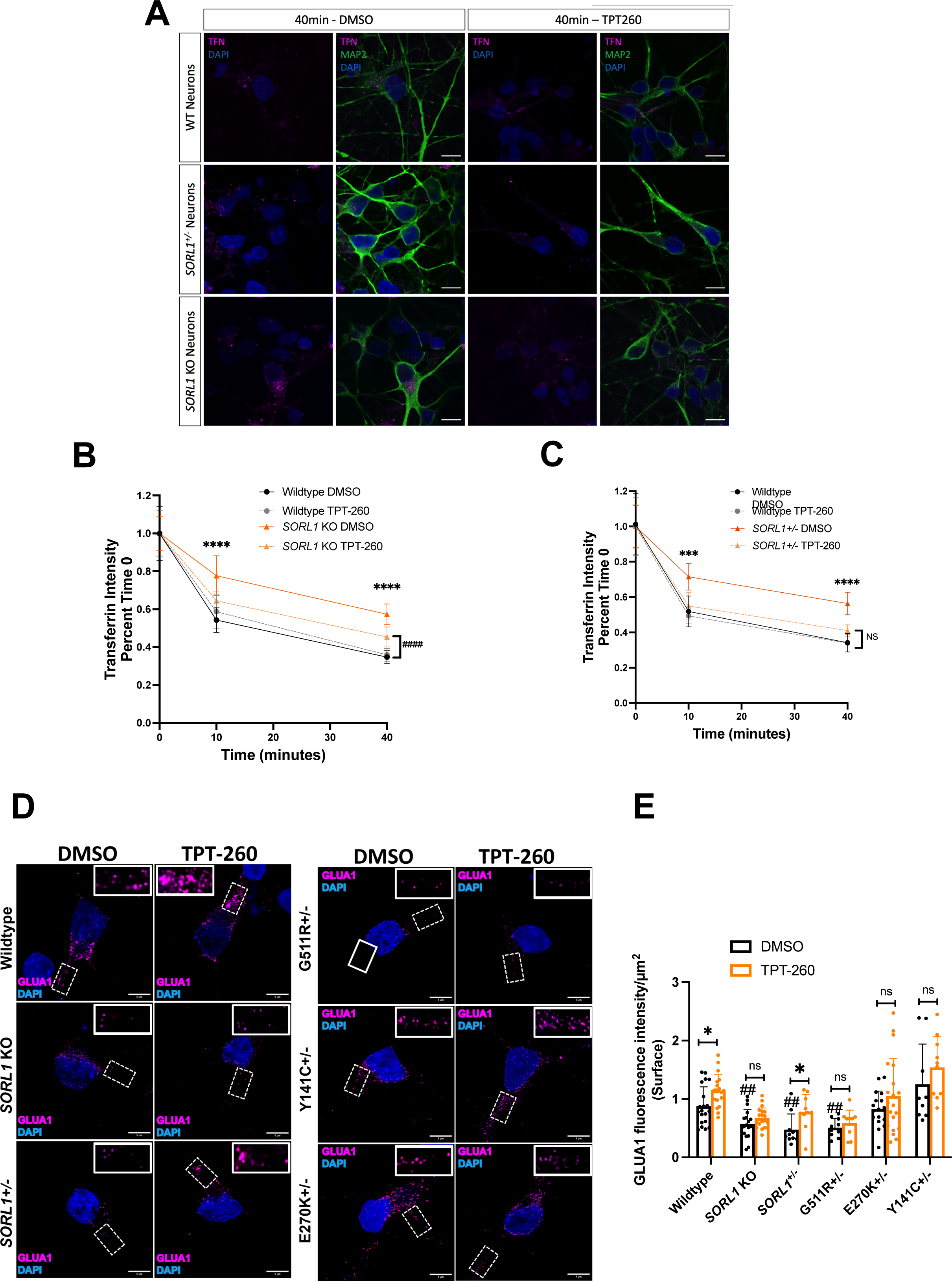
Retromer enhancement with TPT-260 enhances transferrin recycling and surface GLUA1 expression in *SORL1*+/- hiPSC-derived neurons. **(A-C).** The transferrin recycling assay shows a complete rescue of endosomal recycling in *SORL1*_+/-_ neurons treated with TPT-260 (light orange) compared to DMSO (dark orange). **(A)** Representative images showing transferrin recycling in WT, SORL1+/- and SORL1 KO hiPSC-derived neurons. **(B-C)** There is no difference between TPT-260 treated *SORL1_+/-_* neurons and TPT-260 WT neurons (gray lines). Asterisks indicate a statistical difference between WT DMSO and *SORL1*_+/-_ DMSO; NS indicates a non-significant difference between *SORL1*_+/-_ +TPT-260 and WT+TPT-260. TPT-260 only partially rescues endosomal recycling in *SORL1* KO neurons. TPT-260 treated *SORL1* KO neurons (dark orange) do not recycle transferrin as efficiently as WT neurons (black/gray). Asterisks indicate a statistical difference between WT DMSO and *SORL1* KO DMSO neurons. Hashmarks indicate a statistical difference between TPT-260 treated *SORL1* KO and TPT-260 treated WT neurons. Scale bar: 5 μm. Data represented as transferrin intensity normalized to time 0. 10 images per clone per genotype per timepoint were analyzed. 1-3 clones per genotype were analyzed. **(D-E)** Fluorescence intensity of surface GLUA1 measured in DMSO and TPT-260 treated WT, *SORL1* KO and *SORL1* +/- hiPSC-derived neurons. (D-E) Representative images showing decreased fluorescence intensity of surface GLUA1 in DMSO treated *SORL1* KO, *SORL1* +/- and G511R +/- hiPSC-derived neurons as compared to DMSO treated WT hiPSC-derived neurons. Significance indicated by hashmarks. TPT-260 rescues surface GLUA1 expression in SORL1 +/- hiPSC-derived neurons. Quantification done using Image J. 10 images per clone per genotype per timepoint were analyzed. Scale bar: 5μm; 1-3 clones per genotype were analyzed. Each data point on the graph indicates mean GLUA1 fluorescence intensity/image of non-permeabilized cells normalized to mean of permeabilized cells. Data represented as mean ± SD. Normally distributed data was analyzed using parametric two-way ANOVA. Significance was defined as a value of */^#^p < 0.05, **/^##^p < 0.01, ***/^###^p < 0.001, and ****/^####^p < 0.0001. ns= not significant

## Discussion

When considering the development of novel therapeutics for Alzheimer’s disease, it is critical to examine biologically relevant pathways such as protein trafficking through the endo-lysosomal network. In particular, trafficking of APP directly affects its processing into Aβ. Furthermore, indicators of endosomal dysfunction are early cytopathological phenotypes in AD, evident before substantial accumulation of other neuropathologic hallmarks(Cataldo et al., 2000) and is therefore a potentially attractive early therapeutic readout. In order to more fully explore this concept in human neurons, we used previously published and newly generated hiPSC lines that are either deficient in *SORL1* expression (*SORL1* KO or *SORL1*^+/-^) or that harbor AD-associated coding variants in the VPS10 domain of the protein (*SORL1*^Var^) and treated neurons differentiated from these hiPSCs with TPT-260, a small molecule chaperone that stabilized retromer and enhances its function(Mecozzi et al., 2014).

We and others have previously reported that heterozygous and homozygous loss of *SORL1* results in increased secretion of Aβ and enlarged early endosomes(Hung et al., 2021; Knupp et al., 2020). In this study, we also observed increases in secreted Aβ and enlarged endosomes in our *SORL1*^Var^ lines (**Figure 1**). Both secreted Aβ and enlarged early endosome phenotypes seem to be dependent on whether a WT allele of *SORL1* is present, which aligns with previous reports(Dodson et al., 2008; Hung et al., 2021). These data are also consistent with indications that *SORL1* haploinsufficiency is causative for AD(Holstege et al., 2017; Scheltens et al., 2021), and suggests that certain AD-associated variants may result in loss of important *SORL1* functions.

Our data provides the first evidence that a small molecule can correct early endosome enlargement. TPT-260 treatment reduced endosome size in all genotypes, however only in neurons with at least one copy of *SORL1* were endosome sizes fully reduced to WT levels (**Figure 2**). Importantly for therapeutic implications, the retromer chaperone does not appear to affect endosome size in the WT cell lines at the concentrations we tested.

SORLA, is the main adaptor protein for retromer-dependent trafficking of APP and Aβ peptides(Andersen et al., 2005; Caglayan et al., 2014; Fjorback et al., 2012). Therefore, in *SORL1* KO neurons, there is a smaller effect on Aβ secretion after treatment with TPT-260 (**Figure 3).** However, in the more clinically relevant scenario of either *SORL1*^+/-^or *SORL1*^Var^, TPT-260 treatment reduced both Aβ40 and Aβ42 to WT levels. TPT260 treatment lowers phospho-Tau levels on multiple epitopes in this model, highlighting the potential therapeutic relevance of retromer functional enhancement. Our data here (**Figure 3**) provide an independent corroboration of previous work(Young et al., 2018)and also supports what has been seen in tauopathy models(Li et al., 2020; Young et al., 2018). Tau clearance by endo-lysosomal trafficking is an important aspect of maintaining Tau homeostasis and efficient clearance of p-Tau is an important step in preventing pathological aggregation(Tang et al., 2019).

One consistent phenotype we observed across our *SORL1* KO, *SORL1*^+/-^ and *SORL1*^Var^ cell lines were enlarged endosomes, indicative of endosomal traffic jams. Endosomal traffic jams impact multiple arms of the endo-lysosomal network. *SORL1* KO neurons have enlarged lysosomes and impaired lysosomal degradation(Mishra et al., 2022). In this study, we found that only one variant of *SORL1*, G511R, demonstrated enlarged lysosomes and that this phenotype was rescued with TPT-260 (**Figure 4C-D**). When we probed further into the effects of retromer enhancement on lysosomal degradation using the DQ-Red-BSA assay, we observed that TPT-260 enhances lysosomal degradation in *SORL1* KO cells, bringing this function to WT levels (**Figure 4D-E**). Together, these data suggest that retromer enhancement of the retrograde pathway, trafficking from endosome-Golgi-lysosome, can be restored in *SORL1* KO or variant cells.

SORLA is also heavily involved in the endosomal recycling pathway in neurons. We tested this in two ways, first by transferrin recycling and then by analysis of cell surface levels of GLUA1. We observed only a partial rescue of transferrin recycling in *SORL1* KO cells but a full rescue in *SORL1*^+/-^ neurons (**Figure 6A-C**). In neurons, SORLA interacts with a neuron-specific isoform of the retromer complex, VPS26B, to recycle cargo such as GLUA1(Simoes et al., 2021) and this process is impaired in *SORL1* KO neurons(Mishra et al., 2022). Therefore, we analyzed cell surface levels of GLUA1 in *SORL1*^+/-^ and *SORL1*^Var^ neurons. In line with our previous results, we see reduced cell surface staining of GLUA1 in *SORL1* KO, *SORL1*^+/-^, and *SORL1* G511R variant neurons. Interestingly, two variants, E270K and Y141C, do not show reduced cell surface staining and cell surface levels are unchanged after treatment with TPT-260 (**Figure 6D-E**). Treatment with TPT-260 rescues GLUA1 surface expression in *SORL1*^+/-^ neurons, but not in *SORL1* KO or G511R variant cells, suggesting that the G511R variant could be further impeding the function of the WT allele (**Figure 6D-E**). This is the second example of phenotypic differences between variants in our study as only G511R variant cells showed changes in lysosomal morphology (**Figure 4C-D**). There are over 500 identified coding variants in SORL1(Holstege H, 2023). Our study indicates that not all coding variants show similar phenotypes and highlights the need for further studies into the biology and classification of SORL1 variants.

Endosomal traffic jams can lead to mis-localization of important cellular proteins. For example, APP and VPS35 are both increased in early endosomes in *SORL1* KO neurons(Knupp et al., 2020; Mishra et al., 2022), which could contribute to increased amyloidogenic processing of APP to Aβ and alterations in normal retromer-related trafficking. TPT-260 treatment reduced the localization of VPS35 in early endosomes to WT levels in *SORL1* KO cells, showing that this treatment is sufficient to mobilize retromer in these cells (**Figure 5**). In neurons with at least one copy of *SORL1* (*SORL1*^+/-^) neurons, TPT-260 treatment APP co-localization with early endosomes is similar to WT APP/EEA1 co-localization (**Figure 5A-B**). However, because SORLA is a main adaptor protein between retromer and APP, in full *SORL1* KO cells, there is not a full rescue of endosomal APP localization with TPT-260 treatment (**Figure 5C-D**). A large part of amyloidogenic cleavage of APP occurs in the early endosomes, so this finding could explain why Aβ levels in *SORL1*^+/-^ neurons are completely rescued with TPT-260 treatment, while Aβ levels in *SORL1* KO neurons are not.

Our work shows that treatment with a small molecule that stabilizes retromer in *vitro* and enhances retromer-related trafficking can reduce important cellular and neuropathologic phenotypes in human AD neuronal models. Using TPT-260, we broadly show that enhancement of retromer-related trafficking can fully or partially rescue deficits induced by loss of *SORL1*. This includes trafficking of GLUA1 to the neuronal surface, reduction of amyloidogenic APP processing, and increasing lysosomal degradative capacity. In the case of AD, all of those scenarios can likely be considered neuroprotective, thus having them in combination makes a strong case that stabilizing retromer is a valid therapeutic strategy. Although there is one report of a full loss of *SORL1*(Le Guennec et al., 2018), most patients where *SORL1* is a main risk factor for disease development still maintain at least one functional copy of *SORL1* and thus may benefit from enhancement of the SORL1-retromer pathway. However, our studies also show that different *SORL1* variants have varying phenotypic effects in neurons and also differences in the degree of rescue by TPT-260. Therefore, it will be important for future studies to carefully characterize and classify putative pathogenic variants in *SORL1* and specifically targeted therapies could be warranted. We summarize the variants we studied, the phenotypes we observed, the phenotypes that changed with TPT-260 treatment, and other literature on these SORL1 variants in **Figure 7**. This study builds on previous work that has used these chaperones in animals and human cells, suggesting that studies such as this could represent an important pre-clinical step in identifying new therapeutic molecules for AD.

**Figure 7.**
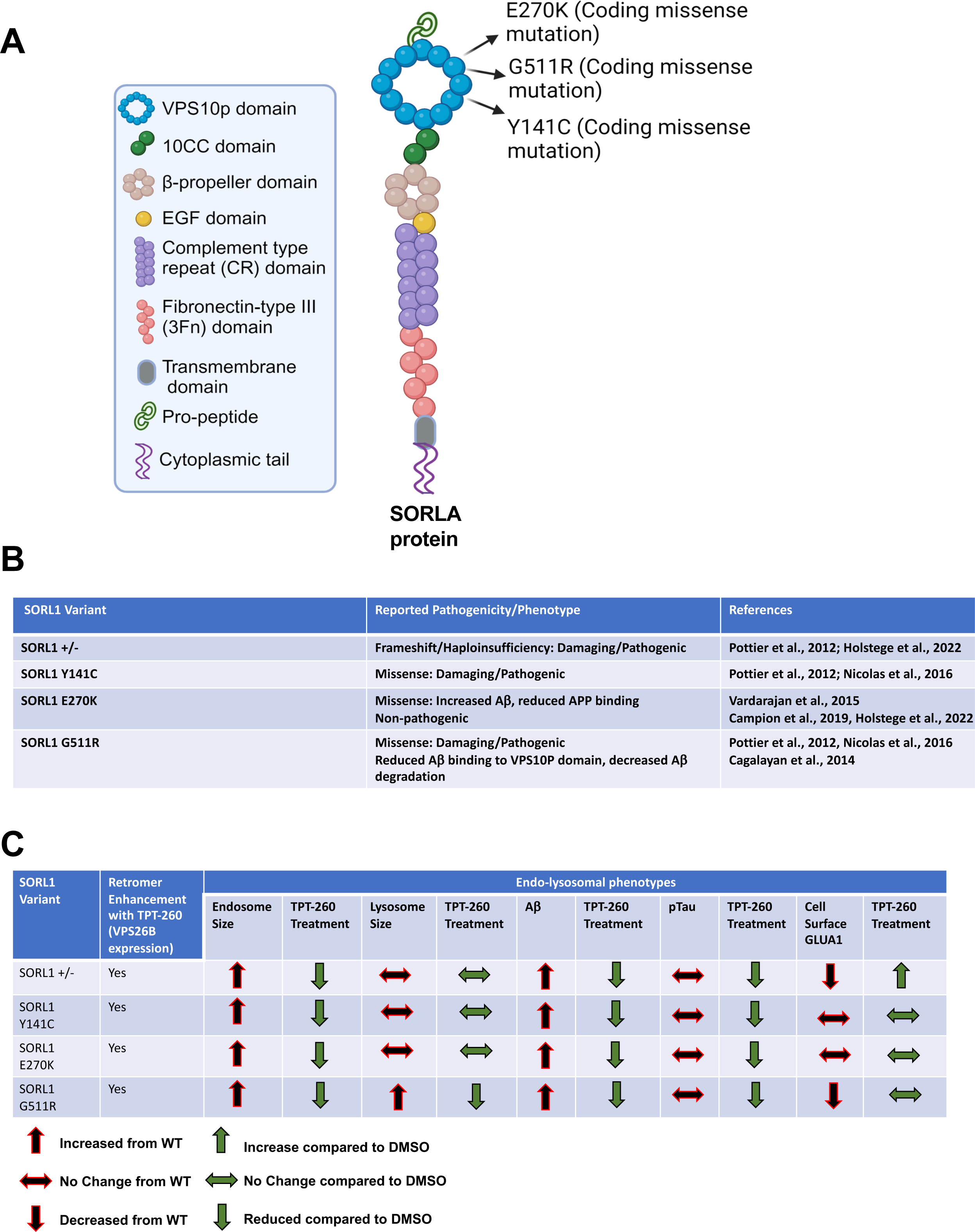
Summary of SORL1 variants and phenotypes studied. **(A)** Schematic of SORLA protein structure depicting location and type of SORL1^var^ mutations used in this study. **(B)** Detailed description of SORL1^var^ mutations and previously studied AD-related pathological phenotypes associated with these mutations **(C)** Schematic showing summary of endo-lysosomal phenotypes observed in SORL1^var^ mutations used in this study and the changes in phenotypes observed with retromer enhancement (TPT-260).

### Limitations to our study

Our study has certain limitations. First, we focused on variants present in the VPS10 domain of *SORL1*(Andersen et al., 2016; Pottier et al., 2012; Vardarajan et al., 2014). Some of these variants have been found in control subjects(Holstege et al., 2017), thus there are other factors in various human genetic backgrounds that need to be considered. We engineered these variants in our well-characterized, male, hiPSC line(Knupp et al., 2020; Young et al., 2015). The genetic background of this cell line also harbors one copy of APOE ε4 and common variants in *SORL1* associated with increased AD risk in candidate-gene based studies(Levy et al., 2007; Rogaeva et al., 2007). Because humans are genetically heterogeneous, we cannot rule out the contribution of these other genomic variants to our phenotypes and acknowledge that this type of treatment may have varying effects across different individuals. This variation may also be evident between male vs. female genetic sex or between different ethnicities. However, rescue of these specific endo-lysosomal phenotypes using retromer-enhancing drugs is still an important observation that may be relevant to both earlier and later-onset forms of AD and corroborates previous work where similar molecules were tested across multiple human genomes(Young et al., 2018). Finally, our study is focused on neuronal cells. We recognize the importance of endosomal trafficking and *SORL1*-retromer related pathways in glia and other cell types relevant to the pathogenesis and/or progression of AD. Future studies will benefit from analyzing endo-lysosomal phenotypes in multiple CNS cell types.

Experimental procedures

### Cell lines

#### CRISPR/Cas9 Genome Editing

All genome editing was completed in the previously published and characterized CV background human induced pluripotent stem cell line(Young et al., 2015), which is male with an APOE e3/e4 genotype(Levy et al., 2007). Genome editing was performed according to published protocols(Knupp et al., 2020; Young et al., 2018). Further details about gene editing are described in the supplemental methods.

#### Neuronal Differentiation

hiPSCs were cultured and differentiated into neurons using dual SMAD inhibition protocols(Shi et al., 2012), modified previously in our laboratory(Mishra et al., 2022). Further details about the neuronal differentiations are described in the supplemental methods.

#### Purification of hiPSC-derived neurons

hiPSC-derived neurons were purified using magnetic bead sorting. Details of the procedure are described in the supplemental methods.

#### DQ-Red BSA assay

Lysosomal proteolytic degradation was evaluated using DQ Red BSA (#D-12051; Thermo Fisher Scientific) following published protocols(Davis et al., 2021; Marwaha and Sharma, 2017; Mishra et al., 2022). More details are described in the supplemental methods.

#### Immunocytochemistry

Details of immunocytochemistry and antibodies used are described in the supplemental procedures.

#### Confocal microscopy, Image processing and colocalization analysis

All microscopy and image processing were performed under blinded conditions. Confocal z stacks were obtained using a Nikon A1R confocal microscope with x63 and x100 plan apochromat oil immersion objectives or a Nikon Yokogawa W1 spinning disk confocal microscope and a 100X plan apochromat oil immersion objective. Image processing was performed using ImageJ software(Schindelin et al., 2012). For endosome analysis, 10-20 fields were analyzed for a total of 10-58 cells. The analysis was focused on the neuronal soma region. To investigate colocalization of APP and VPS35 with early endosomes, hiPSC-derived neurons were co-labeled with either APP or VPS35 and early endosome marker EEA1. A minimum of 10 fields of confocal were captured. Median filtering was used to remove noise from images and Otsu thresholding was applied to all images. Colocalization was quantified using the JACOP plugin in Image J software and presented as Mander’s correlation co-efficient. More details can be found in the supplemental methods.

#### Transferrin recycling assay

To measure recycling pathway function, we utilized transferrin recycling assay as previously described(Mishra et al., 2022; Sakane et al., 2014). More details are described in the supplemental methods.

#### GLUA1 cell surface expression

GLUA1 cell surface expression was quantified as reported previously(Mishra et al., 2022). In brief, Image J software(Schindelin et al., 2012) was used to measure GLUA1 fluorescence intensity in maximum z projections. Cell surface expression was reported as a ratio of non-permeabilized fluorescence to permeabilized fluorescence.

#### Western Blotting

Details of western blotting used in this study are described in the supplemental methods.

#### Measurement of secreted Amyloid Beta 1-40 and 1-42

For all Aβ assays, terminally differentiated neurons were plated at the the same cell number (250,000) cells per well. Aβ peptides were measured using an MSD Aβ V-PLEX assay (Meso Scale Discovery #151200E-2) following manufacturers protocols.

#### Measurement of phosphorylated and total tau by ELISA

For all Tau assays, terminally differentiated neurons were plated at the same cell number (250,000) cells per well. Total and phosphorylated tau protein were measured using a Phospho(Thr231)/Total Tau ELISA plate (Meso Scale Discovery #K15121D-2) following manufacturers protocols.

#### Quantification and statistical analysis

The data here represent, when possible, multiple hiPSC clones. This includes two or three WT clones, two *SORL1*KO clones and one *SORL1*^+/-^ clone. Only one *SORL1*^+/-^ clone was recovered during the gene-editing process. For the *SORL1*^Var^ lines and experiments we analyzed two E270K clones, two G511R clones, and one Y141C clone. Only one Y141C clone was recovered during the gene-editing process. All data represents three independent experiments per clone. Experimental data was tested for normal distributions using the Shapiro-Wilk normality test. Normally distributed data was analyzed using parametric two-tailed unpaired t tests, one-way ANOVA tests, or two-way ANOVA tests. Significance was defined as a value of p > 0.05. All statistical analysis was completed using GraphPad Prism software. Further details of quantification and statistical analysis for all experiments are provided in the supplemental procedures.

## Acknowledgments

This work was supported by NIH grant R01AG062148, a BrightFocus Foundation grant (A2018656S), the WeillNeuroHub and Sponsored Research Agreements from Biogen and Retromer Therapeutics to J.E.Y. A.K. was supported by a NIH training grant (T32 AG052354). S.M is supported by a UW ADRC Development grant. Further support for this work comes from a generous gift from the Ellison Foundation (to UW). We thank all the members of the Young Laboratory as well as Dr. Olav M. Andersen, Dr. Scott A. Small and Dr. Gregory A. Petsko for critical comments, discussions and feedback on this work. We would like to acknowledge the UW SLU Cell Analysis Facility and the Garvey Imaging Core at the UW Institute for Stem Cell and Regenerative Medicine.

## Author Contributions

Conceptualization: J.E.Y, A.K., S.M. Experimental Design: J.E.Y., A.K, and S.M. Experimental performance: A.K., S.M, C.A.W, S.E., R.M, C.K, P.T. Writing-Original draft: A.K. and J.E.Y. Writing-Reviewing and editing: S.M., A.K. and J.E.Y. Funding acquisition: J.E.Y, A.K, and S.M. Supervision: J.E.Y. All authors read and approved the final manuscript.

## Notes

### Competing Interest Statement

The authors have declared no competing interest.

### Summary of Updates

New data, results and discussion

## References

Andersen, O.M., Bogh, N., Landau, A.M., Ploen, G.G., Jensen, A.M.G., Monti, G., Ulhoi, B.P., Nyengaard, J.R., Jacobsen, K.R., Jorgensen, M.M., et al. (2022). A genetically modified minipig model for Alzheimer’s disease with SORL1 haploinsufficiency. Cell Rep Med 3, 100740.

Andersen, O.M., Monti, G, Jensen A.M.G., de Waal, M., Hulsman, M., Olsen J.G., Holstege, H. (2023). Relying on the relationship with known disease-causing variants in homologus proteins to predict pathogenicity of SORL1 variants in Alzheimer’s disease. BioRxiv doi10.1101/2023.02.27.524103.

Andersen, O.M., Reiche, J., Schmidt, V., Gotthardt, M., Spoelgen, R., Behlke, J., von Arnim, C.A., Breiderhoff, T., Jansen, P., Wu, X., et al. (2005). Neuronal sorting protein-related receptor sorLA/LR11 regulates processing of the amyloid precursor protein. Proc Natl Acad Sci U S A 102, 13461–13466.

Andersen, O.M., Rudolph, I.M., and Willnow, T.E. (2016). Risk factor SORL1: from genetic association to functional validation in Alzheimer’s disease. Acta Neuropathol 132, 653–665.

Caglayan, S., Takagi-Niidome, S., Liao, F., Carlo, A.S., Schmidt, V., Burgert, T., Kitago, Y., Fuchtbauer, E.M., Fuchtbauer, A., Holtzman, D.M., et al. (2014). Lysosomal sorting of amyloid-beta by the SORLA receptor is impaired by a familial Alzheimer’s disease mutation. Sci Transl Med 6, 223ra220.

Cataldo, A.M., Hamilton, D.J., Barnett, J.L., Paskevich, P.A., and Nixon, R.A. (1996). Properties of the endosomal-lysosomal system in the human central nervous system: disturbances mark most neurons in populations at risk to degenerate in Alzheimer’s disease. J Neurosci 16, 186–199.

Cataldo, A.M., Petanceska, S., Terio, N.B., Peterhoff, C.M., Durham, R., Mercken, M., Mehta, P.D., Buxbaum, J., Haroutunian, V., and Nixon, R.A. (2004). Abeta localization in abnormal endosomes: association with earliest Abeta elevations in AD and Down syndrome. Neurobiol Aging 25, 1263–1272.

Cataldo, A.M., Peterhoff, C.M., Troncoso, J.C., Gomez-Isla, T., Hyman, B.T., and Nixon, R.A. (2000). Endocytic pathway abnormalities precede amyloid beta deposition in sporadic Alzheimer’s disease and Down syndrome: differential effects of APOE genotype and presenilin mutations. Am J Pathol 157, 277–286.

Chu, J., and Pratico, D. (2017). The retromer complex system in a transgenic mouse model of AD: influence of age. Neurobiol Aging 52, 32–38.

Das, U., Wang, L., Ganguly, A., Saikia, J.M., Wagner, S.L., Koo, E.H., and Roy, S. (2016). Visualizing APP and BACE-1 approximation in neurons yields insight into the amyloidogenic pathway. Nat Neurosci 19, 55–64.

Davis, S.E., Roth, J.R., Aljabi, Q., Hakim, A.R., Savell, K.E., Day, J.J., and Arrant, A.E. (2021). Delivering progranulin to neuronal lysosomes protects against excitotoxicity. J Biol Chem 297, 100993.

Dodson, S.E., Andersen, O.M., Karmali, V., Fritz, J.J., Cheng, D., Peng, J., Levey, A.I., Willnow, T.E., and Lah, J.J. (2008). Loss of LR11/SORLA enhances early pathology in a mouse model of amyloidosis: evidence for a proximal role in Alzheimer’s disease. J Neurosci 28, 12877–12886.

Fazeli, E., Child, D.D., Bucks, S.A., Stovarsky, M., Edwards, G., Yu, C.E., Latimer, C., Kitago, Y., Bird, T., Andersen, O.M., et al. (2023). A familial missense variant in the AD gene SORL1 impairs its maturation and endosomal sorting. bioRxiv.

Fjorback, A.W., Seaman, M., Gustafsen, C., Mehmedbasic, A., Gokool, S., Wu, C., Militz, D., Schmidt, V., Madsen, P., Nyengaard, J.R., et al. (2012). Retromer binds the FANSHY sorting motif in SorLA to regulate amyloid precursor protein sorting and processing. The Journal of neuroscience : the official journal of the Society for Neuroscience 32, 1467–1480.

Holstege H, d.M., tesi N, vanderLee SJ, ADESconsortium, ADSPconsortium, StEP-ADconsortium, Knight-ADRC, UCSF/NYGC/UAB, Vogel M, vanSpaendonk R, Hulsman M, Andersen OM. (2023). Effect of prioritized SORL1 missense variants supports clinical consideration for familial Alzheimer’s Disease. medRxiv 10.1101/2023.07.13.23292622.

Holstege, H., Hulsman, M., Charbonnier, C., Grenier-Boley, B., Quenez, O., Grozeva, D., van Rooij, J.G.J., Sims, R., Ahmad, S., Amin, N., et al. (2022). Exome sequencing identifies rare damaging variants in ATP8B4 and ABCA1 as risk factors for Alzheimer’s disease. Nat Genet 54, 1786–1794.

Holstege, H., van der Lee, S.J., Hulsman, M., Wong, T.H., van Rooij, J.G., Weiss, M., Louwersheimer, E., Wolters, F.J., Amin, N., Uitterlinden, A.G., et al. (2017). Characterization of pathogenic SORL1 genetic variants for association with Alzheimer’s disease: a clinical interpretation strategy. Eur J Hum Genet 25, 973–981.

Hung, C., Tuck, E., Stubbs, V., van der Lee, S.J., Aalfs, C., van Spaendonk, R., Scheltens, P., Hardy, J., Holstege, H., and Livesey, F.J. (2021). SORL1 deficiency in human excitatory neurons causes APP-dependent defects in the endolysosome-autophagy network. Cell Rep 35, 109259.

Jensen AM, R.J., Fojtik P, Monti G, Lunding M, Vochyanova S, Pospislova V, vanderLee SJ, VanDongen J, Bossaerts L, VanBroeckhoven C, Dols O, Lleo A, Benussi L, Ghidoni R, Hulsman M, Sleegers K, Bohaciakova D, Holstege H, Andersen O. (2023). The SORL1 p.Y1816C varaint causes impaired endosomal dimerization and autosomal dominant Alzheimer’s disease. MedRxiv 10.1101/2023.07.09.23292253.

Karch, C.M., and Goate, A.M. (2015). Alzheimer’s disease risk genes and mechanisms of disease pathogenesis. Biol Psychiatry 77, 43–51.

Knupp, A., Mishra, S., Martinez, R., Braggin, J.E., Szabo, M., Kinoshita, C., Hailey, D.W., Small, S.A., Jayadev, S., and Young, J.E. (2020). Depletion of the AD Risk Gene SORL1 Selectively Impairs Neuronal Endosomal Traffic Independent of Amyloidogenic APP Processing. Cell reports 31, 107719.

Le Guennec, K., Tubeuf, H., Hannequin, D., Wallon, D., Quenez, O., Rousseau, S., Richard, A.C., Deleuze, J.F., Boland, A., Frebourg, T., et al. (2018). Biallelic Loss of Function of SORL1 in an Early Onset Alzheimer’s Disease Patient. J Alzheimers Dis 62, 821–831.

Lee, H., Aylward, A.J., Pearse, R.V., 2nd, Lish, A.M., Hsieh, Y.C., Augur, Z.M., Benoit, C.R., Chou, V., Knupp, A., Pan, C., et al. (2023). Cell-type-specific regulation of APOE and CLU levels in human neurons by the Alzheimer’s disease risk gene SORL1. Cell reports, 112994.

Levy, S., Sutton, G., Ng, P.C., Feuk, L., Halpern, A.L., Walenz, B.P., Axelrod, N., Huang, J., Kirkness, E.F., Denisov, G., et al. (2007). The diploid genome sequence of an individual human. PLoS Biol 5, e254.

Li, J.G., Chiu, J., Ramanjulu, M., Blass, B.E., and Pratico, D. (2020). A pharmacological chaperone improves memory by reducing Abeta and tau neuropathology in a mouse model with plaques and tangles. Mol Neurodegener 15, 1.

Marwaha, R., and Sharma, M. (2017). DQ-Red BSA Trafficking Assay in Cultured Cells to Assess Cargo Delivery to Lysosomes. Bio Protoc 7.

Mecozzi, V.J., Berman, D.E., Simoes, S., Vetanovetz, C., Awal, M.R., Patel, V.M., Schneider, R.T., Petsko, G.A., Ringe, D., and Small, S.A. (2014). Pharmacological chaperones stabilize retromer to limit APP processing. Nature chemical biology 10, 443–449.

Mishra, S., Knupp, A., Szabo, M.P., Williams, C.A., Kinoshita, C., Hailey, D.W., Wang, Y., Andersen, O.M., and Young, J.E. (2022). The Alzheimer’s gene SORL1 is a regulator of endosomal traffic and recycling in human neurons. Cell Mol Life Sci 79, 162.

Muzio, L., Sirtori, R., Gornati, D., Eleuteri, S., Fossaghi, A., Brancaccio, D., Manzoni, L., Ottoboni, L., Feo, L., Quattrini, A., et al. (2020). Retromer stabilization results in neuroprotection in a model of Amyotrophic Lateral Sclerosis. Nat Commun 11, 3848.

Ouellette, S.P., and Carabeo, R.A. (2010). A Functional Slow Recycling Pathway of Transferrin is Required for Growth of Chlamydia. Front Microbiol 1, 112.

Pottier, C., Hannequin, D., Coutant, S., Rovelet-Lecrux, A., Wallon, D., Rousseau, S., Legallic, S., Paquet, C., Bombois, S., Pariente, J., et al. (2012). High frequency of potentially pathogenic SORL1 mutations in autosomal dominant early-onset Alzheimer disease. Mol Psychiatry 17, 875–879.

Rogaeva, E., Meng, Y., Lee, J.H., Gu, Y., Kawarai, T., Zou, F., Katayama, T., Baldwin, C.T., Cheng, R., Hasegawa, H., et al. (2007). The neuronal sortilin-related receptor SORL1 is genetically associated with Alzheimer disease. Nat Genet 39, 168–177.

Sakane, H., Horii, Y., Nogami, S., Kawano, Y., Kaneko-Kawano, T., and Shirataki, H. (2014). alpha-Taxilin interacts with sorting nexin 4 and participates in the recycling pathway of transferrin receptor. PLoS One 9, e93509.

Scheltens, P., De Strooper, B., Kivipelto, M., Holstege, H., Chetelat, G., Teunissen, C.E., Cummings, J., and van der Flier, W.M. (2021). Alzheimer’s disease. Lancet 397, 1577–1590.

Schindelin, J., Arganda-Carreras, I., Frise, E., Kaynig, V., Longair, M., Pietzsch, T., Preibisch, S., Rueden, C., Saalfeld, S., Schmid, B., et al. (2012). Fiji: an open-source platform for biological-image analysis. Nat Methods 9, 676–682.

Shi, Y., Kirwan, P., Smith, J., Robinson, H.P., and Livesey, F.J. (2012). Human cerebral cortex development from pluripotent stem cells to functional excitatory synapses. Nat Neurosci 15, 477–486, S471.

Simoes, S., Guo, J., Buitrago, L., Qureshi, Y.H., Feng, X., Kothiya, M., Cortes, E., Patel, V., Kannan, S., Kim, Y.H., et al. (2021). Alzheimer’s vulnerable brain region relies on a distinct retromer core dedicated to endosomal recycling. Cell Rep 37, 110182.

Sonnichsen, B., De Renzis, S., Nielsen, E., Rietdorf, J., and Zerial, M. (2000). Distinct membrane domains on endosomes in the recycling pathway visualized by multicolor imaging of Rab4, Rab5, and Rab11. J Cell Biol 149, 901–914.

Tan, J.Z.A., and Gleeson, P.A. (2019). The role of membrane trafficking in the processing of amyloid precursor protein and production of amyloid peptides in Alzheimer’s disease. Biochim Biophys Acta Biomembr 1861, 697–712.

Tang, M., Harrison, J., Deaton, C.A., and Johnson, G.V.W. (2019). Tau Clearance Mechanisms. Adv Exp Med Biol 1184, 57–68.

Toh, W.H., Chia, P.Z.C., Hossain, M.I., and Gleeson, P.A. (2018). GGA1 regulates signal-dependent sorting of BACE1 to recycling endosomes, which moderates Abeta production. Mol Biol Cell 29, 191–208.

Vagnozzi, A.N., Li, J.G., Chiu, J., Razmpour, R., Warfield, R., Ramirez, S.H., and Pratico, D. (2021). VPS35 regulates tau phosphorylation and neuropathology in tauopathy. Molecular psychiatry 26, 6992–7005.

Vardarajan, B.N., Zhang, Y., Lee, J.H., Cheng, R., Bohm, C., Ghani, M., Reitz, C., Reyes-Dumeyer, D., Shen, Y., Rogaeva, E., et al. (2014). Coding mutations in SORL1 and Alzheimer’s disease. Ann Neurol.

Young, J.E., Boulanger-Weill, J., Williams, D.A., Woodruff, G., Buen, F., Revilla, A.C., Herrera, C., Israel, M.A., Yuan, S.H., Edland, S.D., et al. (2015). Elucidating Molecular Phenotypes Caused by the SORL1 Alzheimer’s Disease Genetic Risk Factor Using Human Induced Pluripotent Stem Cells. Cell Stem Cell 16, 373–385.

Young, J.E., Fong, L.K., Frankowski, H., Petsko, G.A., Small, S.A., and Goldstein, L.S.B. (2018). Stabilizing the Retromer Complex in a Human Stem Cell Model of Alzheimer’s Disease Reduces TAU Phosphorylation Independently of Amyloid Precursor Protein. Stem Cell Reports 10, 1046–1058.

