## Supplemental Figures and Methods for "Pharmacologic Stabilization of Retromer Rescues Endosomal Pathology Induced by Defects in the Alzheimer’s gene *SORL1*"

Supplemental Figure 1

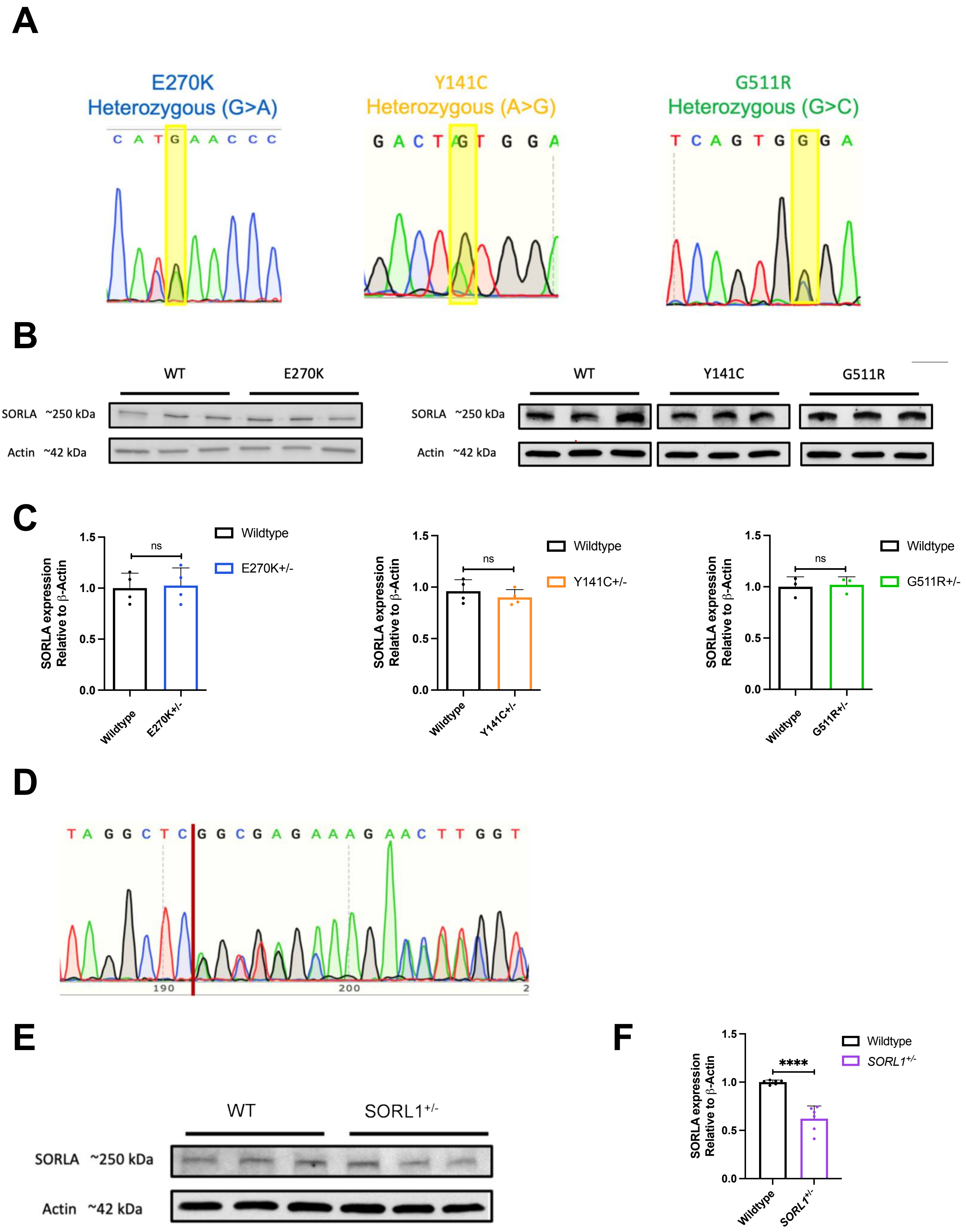

Supplemental Figure 2

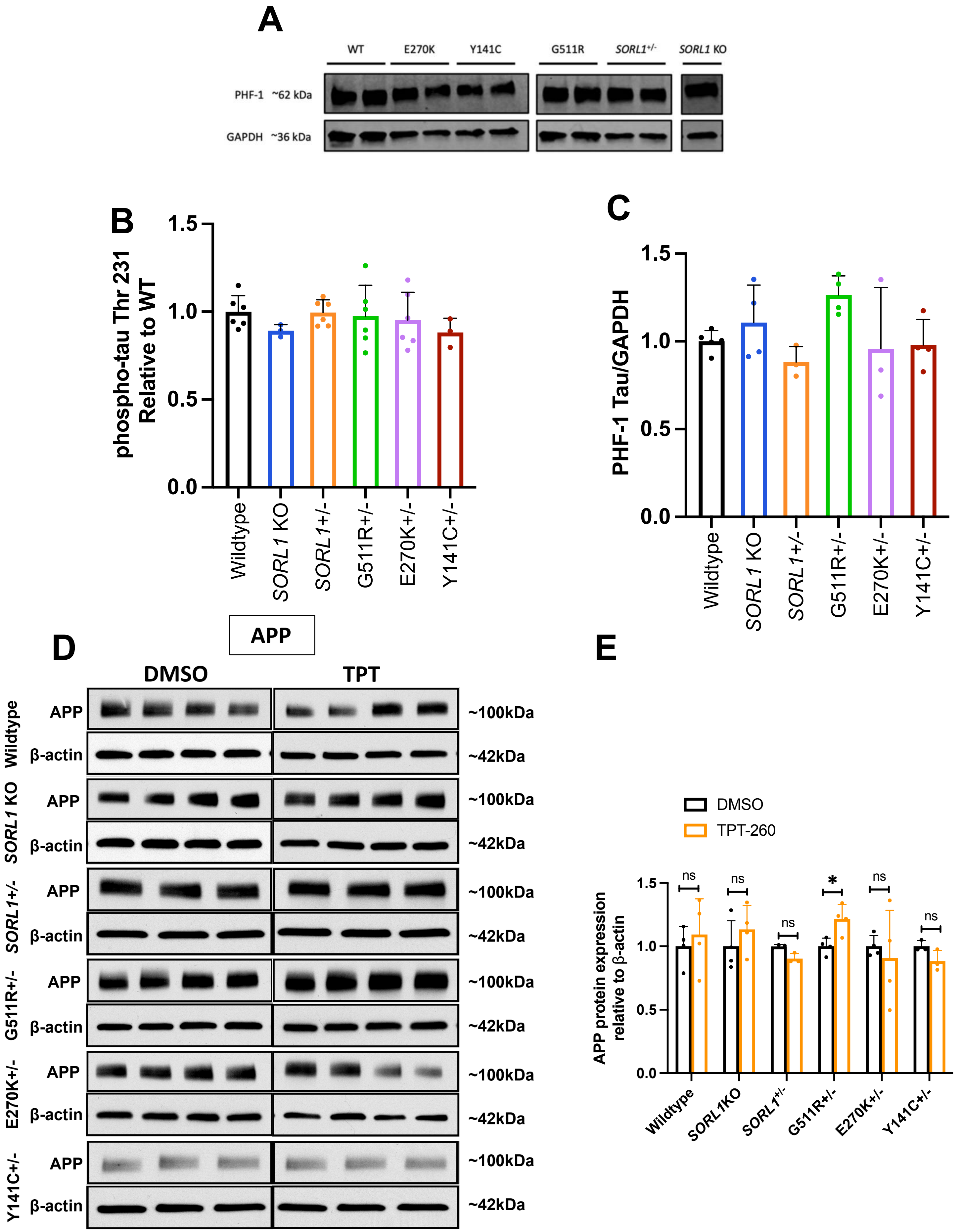

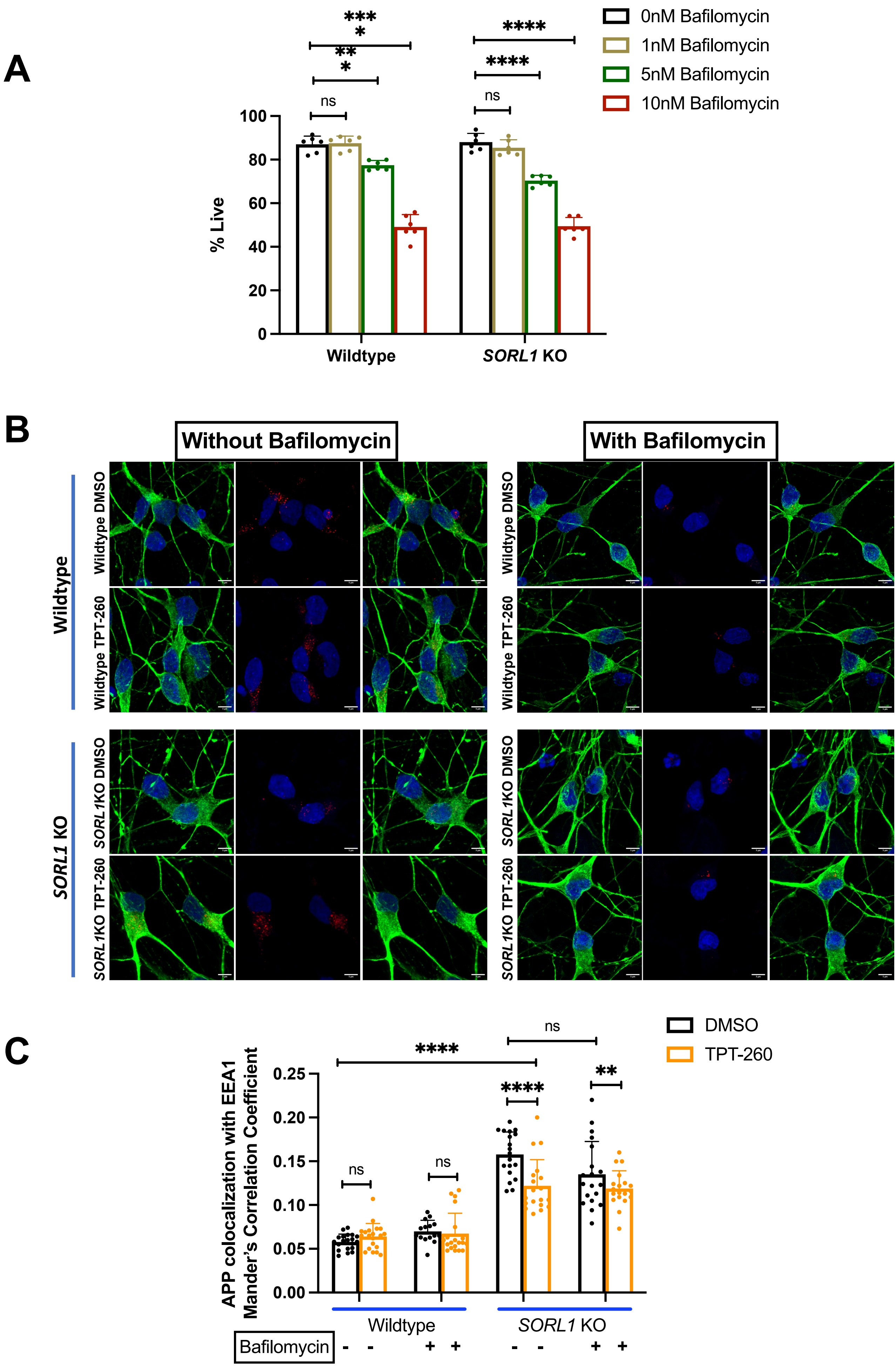

A
0nM Bafilomycin
1nM Bafilomycin
5nM Bafilomycin
10nM Bafilomycin
% Live
0
20
40
60
80
100
ns
***
*
**
*
****
****
ns
Wildtype
SORL1 KO
B
Without Bafilomycin
With Bafilomycin
Wildtype
SORL1 KO
Wildtype DMSO
Wildtype TPT-260
SORL1KO DMSO
SORL1KO TPT-260
C
DMSO
TPT-260
APP colocalization with EEA1
Mander's Correlation Coefficient
0.00
0.05
0.10
0.15
0.20
0.25
ns
ns
****
****
**
ns
Wildtype
SORL1 KO
Bafilomycin
- - + + - - + +

**Supplementary figures**

**Figure S1. Characterization of *SORL1*$^{Var}$ and *SORL1*$^{+/-}$ cell lines.**

**(A)** Sanger sequencing showing heterozygous base-pair change for each *SORL1*$^{Var}$ hiPSC line. Note: the A/T change 5’ to the G>A edit is an endogenous synonymous SNP in the genetic background of the parental hiPSC lines. **(B)** Representative Western blots showing that *SORL1*$^{Var}$ hiPSCs have equivalent levels of SORLA protein expression to WT**. (C)** Quantification of Western blots; E270K analysis: 2 WT and 2 E270K clones, 2 independent replicates per clone;Y141C analysis: 2 WT clones and 1 Y141C clone, 2 independent replicates per clone for WT and 4 independent replicates per clone for Y141C; G511R analysis: 1 WT clone, 3 independent replicates per clone. 1 G511R clone, 3 independent replicates per clone. **(D)** Sanger sequencing showing heterozygous insertion/deletion (indel) leading to *SORL1*$^{+/-}$ hiPSC line. **(E)** Representative Western blots showing that *SORL1*$^{+/-}$ hiPSCs have less SORLA protein expression than WT hiPSCs. **(F)** Quantification of Western blots; *SORL1*$^{+/-}$ analysis: 2 WT clones, 3 independent replicates per clone. 1 *SORL1*$^{+/-}$ clone, 6 independent replicates per clone. Data represented as mean ± SD. Normally distributed data was analyzed using parametric two-way ANOVA. 1-3 clones were analyzed per genotype. Significance was defined as a value of *$p < 0.05$, **$p < 0.01$, ***$p < 0.001$, and ****$p < 0.0001$. ns= not significant

**Figure S2. Phospho-tau levels in *SORL1*KO, *SORL1*$^{Var}$ and *SORL1*$^{+/-}$ neurons. Related to Figure 1-3.**

**(A-C)** No significant change in phospho-tau at two epitopes was detected in *SORL1* KO, *SORL1*$^{Var}$, or *SORL1*$^{+/-}$ neurons. **(A)** Representative Western blot of PHF-1 phospho-tau epitope. **(B)** Quantification of Western blot; WT - 2 clones, 2 independent replicates for clone 1 (A7) and 3 independent replicates for clone 2 (A6); E270K : 2 clones, 2 independent replicates per clone; Y141C : 1 clone, 3 independent replicates per clone; G511R : 2 clones, 2 independent replicates per clone; *SORL1*$^{+/-}$ : 1 clone, 3 independent replicates per clone; *SORL1* KO : 2 clones, 2 independent replicates per clone **(C)** ELISA assay of Thr 231 phospho-Tau epitope; WT : 2 clones, 3 independent replicates per clone; E270K : 2 clones, 3 independent replicates per clone; Y141C : 1 clone, 3 independent replicates per clone ; G511R: 2 clones, 3 independent replicates per clone; *SORL1*$^{+/-}$ : 1 clone, 3 independent replicates per clone; *SORL1* KO : 2 clones, 3 independent replicates per clone. **(D-E)** Western blot showing protein expression of full-length Amyloid precursor protein (APP) in DMSO and TPT-260 treated WT, SORL1 KO, SORL1 +/- and SORL1var hiPSC-derived neurons. Quantification done using Image J software. For this experiment, N=4; 2 clones/genotype of WT, *SORL1* KO, G511R+/- and E270K+/- and 2 replicates/clone were used. For Y141C+/- and *SORL1*+/- cell lines, 1 clone/genotype and 3 replicates/clone were used. Data represented as mean ± SD. Normally distributed data was analyzed using parametric two-way ANOVA. 1-3 clones were analyzed per genotype. Significance was defined as a value of *p < 0.05, **p < 0.01, ***p < 0.001, and ****p < 0.0001. ns= not significant

**Figure S3. Retromer does not alter APP trafficking through the lysosome pathway.**

**(A)** Cytotoxicity of lysosome function inhibitor, Bafilomycin measured using trypan blue assay. WT and *SORL1* KO hiPSC-derived neurons were treated with 0-10nM Bafilomycin and cytotoxicity was measured by calculating % live cells using trypan blue. 1nM was determined as least toxic and 10nM was determined as most toxic in both cell lines. 1nM Bafilomycin was used for future experiments. **(B)** Representative images of DMSO and TPT-260 treated WT and *SORL1* KO hiPSC-derived neurons treated with or without bafilomycin(1nM) showing decreased degradation of DQ-BSA in all the bafilomycin treated cells irrespective of DMSO or TPT-260 treatment, confirming that 1nM bafilomycin sufficiently inhibits lysosomal degradation in all the cell lines used for this experiment. **(C)** Quantification of colocalization of APP with EEA1 in WT and SORL1 KO hiPSC-derived neurons showing no effect on colocalization upon bafilomycin treatment in both DMSO and TPT-260 treated cells. 10 images per clone per genotype per timepoint were analyzed. 2 clones per genotype were analyzed. Each data point on the graph indicates Mander’s correlation co-efficient measured per image. Data represented as mean ± SD. Normally distributed data was analyzed using parametric two-way ANOVA. Significance was defined as a value of *$p < 0.05$, **$p < 0.01$, ***$p < 0.001$, and ****$p < 0.0001$. ns= not significant

**Supplemental Experimental Procedures**

**Cell lines (CRISPR/Cas9 Genome Editing)**
The Zhang Lab CRISPR Design website (crispr.mit.edu) was used to generate guide RNAs (gRNAs) with minimal off-target effects. gRNAs were cloned into the px458 vector that also expresses GFP and the Cas-9 nuclease. hiPSCs were electroporated with plasmids, sorted by flow cytometry for GFP expression, and plated at a clonal density of ~1x10^4 cells per 10cm plate. After approximately two weeks of growth, colonies were picked and split into identical sets. Further details of the cell lines are described in the supplemental experimental procedures, and Tables S1. One set was analyzed by Sanger sequencing and one set was expanded to generate isogenic cell lines.

A total of 8 previously unpublished clones were chosen for experiments contained in this publication: 2 WT clones, 2 clones containing the heterozygous *SORL1* E270K variant (Figure S1), 1 clone containing the heterozygous *SORL1* Y141C variant (Figure S1), 2 clones containing the heterozygous *SORL1* G511R variant (Figure S1), and 1 clone containing a deletion resulting in a heterozygous *SORL1* KO (*SORL1* $^{+/-}$ ) (Figure S2). Also included in this publication are four previously published clones including 2 WT clones and 2 homozygous *SORL1* KO clones (*SORL1* KO)(1, 2). All clones were shown to have normal karyotypes and verified to be free of mycoplasma (MycoAlert).

**CRISPR/Cas9 gRNA, ssODN, and Primer Sequences**

The CRISPR/Cas9 reagents used to generate the E270K cell lines are the same as what was used to generate the previously published *SORL1* KO hiPSC lines(1). For the *SORL1* KO lines, clones were chosen that incorporated indels rather than the ssODN sequence.

E270K gRNA: ATTGAACGACATGAACCCTC
E270K ssODN:
GGGAATTGATCCCTATGACAAACCAAATACCATCTACATTGAACGACATGAACCCTCTGGCTACTCCA
CGTCTTCCGAAGTACAGATTTCTTCCAGTCCCGGGAAAACCAGGAAG
E270K Forward primer: ctctatcctgagtcaaggagtaac
E270K Reverse primer: ccttccaattcctgtgtatgc
PCR amplifies 458 bp sequence

Y141C gRNA: GTACGTGTCTTACGACTA
Y141C ssODN:
GAAAGATCTTTCTGCCAGTTTCTCACCAACTCTTTCTTTTTCATCTCCTTTTCTCTGTATTCCAGGTGT
ACGTGTCTTACGACTGTGGAAAATCATTCAAGAAAATTTCAGACAAGTTAAACTTTGGCTTGGGAAAT
AGGAGTGAAGCTG
Y141C Forward primer: gtggcaggtgcctgtaatcc
Y141C Reverse primer: cacagagagcgccatctcc
PCR amplifies 445 bp sequence

G511R gRNA: CTCTTGCATTTTAGGCTCAG
G511R ssODN:
CTGACATATTCTTGAAATTAAAAATAATTATTTCTCTTGCATTTTAGGCTCAGTGCGAAAGAACTTGGC
TAGCAAGACAAACGTGTACATCTCTAGCAGTGCTGGAGCCAGGTGGCG
G511R Forward primer: cgccactggtaagtgtgcttgc
G511R Reverse primer: ctggcattactggtctctgccatg
PCR amplifies 413 bp sequence

*SORL1*$^{+/-}$: The *SORL1*$^{+/-}$ line was generated with the same gRNA as the *SORL1* G511R line. The PCR primers and amplicon are the same as for G511R.

**Neuronal Differentiation**
Cortical neurons were differentiated from hiPSCs using the dual-SMAD inhibition technique. hiPSCs were plated on 1:20 Matrigel-coated (Growth factor reduced basement membrane matrix; # 356231; Corning) 6-

well plates at a density of 3.5 million cells per well. Cells were fed with Basal Neural Maintenance Media (BNMM) (1:1 DMEM/F12 (#11039047 Life Technologies) + glutamine media/neurobasal media (#21103049, Gibco), 0.5% N2 supplement (# 17502-048; Thermo Fisher Scientific,) 1% B27 supplement (# 17504-044; Thermo Fisher Scientific), 0.5% GlutaMax (# 35050061; Thermo Fisher Scientific), 0.5% insulin-transferrin-selenium (#41400045; Thermo Fisher Scientific), 0.5% NEAA (# 11140050; Thermo Fisher Scientific), 0.2% β-mercaptoethanol (#21985023, Life Technologies)) supplemented with 10 μM SB-431542 and 0.5 μM LDN-193189 (#1062443, Biogems) for seven days. On day 8, cells were incubated with Versene (#15640066, Gibco), dissociated with cell scrapers, and passaged 1:3. From days 9-13, cells were fed daily with BNMM containing no supplements. On day 13, media was switched to BNMM containing 20 ng/mL FGF (R&D Systems, Minneapolis, MN). On day 16, cells were passaged 1:3. Cells were fed daily until approximately day 23, when cells were FACS sorted to enrich a stable population of CD184/CD24 (#557145/561646 BD Pharmingen) positive, CD44/CD271 (#555479/557196 BD Pharmingen) negative neural progenitor cells(3). After sorting, cells were expanded for cortical neuronal differentiation. Neural progenitor cells were plated on Matrigel at a density of 5 million cells per 10cm plate. Media was switched to BNMM supplemented with 0.02 μg/mL brain-derived neurotrophic factor (#450-02 PeproTech) + 0.02 μg/mL glial-cell-derived neurotrophic factor (#450-10 PeproTech) + 0.5 mM dbcAMP (#D0260 Sigma Aldrich). Cells were fed twice a week for three weeks. After three weeks, neurons were sorted by magnetic activated techniques to enrich the population of CD184/CD44/CD271 negative cells and plated out for experiments. All cell culture was maintained at 37C and 5% $CO_2$.

**Purification of hiPSC-derived neurons**
Following 3 weeks of differentiation, neurons were dissociated with accutase and resuspended in Magnet Activated Cell Sorting (MACS) buffer (PBS + 0.5% bovine serum albumin [Sigma Aldrich, St Louis, MO] + 2 mM ethylenediaminetetraacetic acid [Thermo Fisher Scientific, Waltham, MA]). Following a modification of(3) cells were incubated with PE-conjugated mouse anti-Human CD44 and mouse anti-Human CD184 antibodies (BD Biosciences, San Jose, CA) at a concentration of 5 μl/10 million cells. Following antibody incubation, cells were washed with MACS buffer and incubated with anti-PE magnetic beads (BD Biosciences, San Jose, CA) at a concentration of 25 μl/10 million cells. Bead-antibody complexes were pulled down using a rare-earth magnet, supernatants were selected, washed, and plated at an appropriate density.

**DQ Red BSA assay**
Lysosomal proteolytic degradation was evaluated using DQ Red BSA (#D-12051; Thermo Fisher Scientific), a fluorogenic substrate for lysosomal proteases, that generates fluorescence only when enzymatically cleaved in intracellular lysosomal compartments. hiPSC-derived neurons were seeded at a density of 400,000 cells/well of a Matrigel coated 48-well plate. After 24 h, cells were washed once with DPBS, treated with complete media containing either 10 μg/ml DQ Red BSA or vehicle (PBS) and incubated for 24 h at 37 °C in a 5% CO2 incubator as described in(4) and(5). At the end of 24 h, cells were washed with PBS, fixed with 4% PFA and immunocytochemistry was performed as described below in Supplemental Experimental Procedures. Cells were imaged using a Leica SP8 confocal microscope and all image processing was completed with ImageJ software. Cell bodies were identified by MAP2 labeling, and fluorescence intensity of DQ Red BSA was measured in regions of the images containing the MAP2 label. Data was presented as fluorescence intensity of DQ-BSA normalized to cell area. Bafilomycin (#B1793, Sigma) was used for inhibition of lysosomal degradation at a concentration of 1nM.

**Transferrin recycling assay**
Purified neurons were seeded at 400,000 cells/well of a 24-well plate containing Matrigel coated 12 mm glass coverslips/well. After 5 DIV, cells were washed once with DMEM-F12 medium and incubated with starving medium (DMEM-F12 medium + 25 mM HEPES + 0.5% BSA) for 30 min at 37 °C in a 5% CO2 incubator to remove any residual transferrin. Thereafter, cells were pulsed with either 100 μg/ml transferrin from human serum conjugated with Alexa Fluor™ 647(#T23366; Thermo Fisher Scientific) or vehicle (PBS) in ‘starving medium’. At the end of 10 min, cells were washed twice with ice-cold PBS to remove any

external transferrin and stop internalization of transferrin and washed once with acid stripping buffer (25 mM citric acid + 24.5 mM sodium citrate + 280 mM sucrose + 0.01 mM Deferoxamine) to remove any membrane bound transferrin. Next, cells were either fixed in 4% PFA or 'Chase medium' (DMEM-F12 + 50 μM Deferoxamine + 20 mM HEPES + 500 μg/ml Holo-transferrin) was added for different time points. Immunocytochemistry was done using MAP2 antibody to label neurons, confocal images were captured using Leica SP8 confocal microscope under blinded conditions. Fluorescence intensity of transferrin was measured using ImageJ software and presented as transferrin intensity normalized to cell area.

**Western blotting**

Cell lysates were run on 4%–20% Mini-PROTEAN TGX Precast Protein Gels (#4561096; Biorad) or 16.5% Criterion Tris-Tricine Gel (#3450063; BioRad) and transferred to PVDF membranes. Membranes were probed with antibodies described in the Key resources table. Imaging was performed with a BioRad ChemiDoc system and quantification was performed using ImageJ software. For Western blot analysis, hiPSC-neurons were washed with PBS and harvested in RIPA buffer containing 1X protease and 1X phosphatase inhibitors. Total protein concentration was quantified using Pierce BCA assay kit (#23225; Thermo Fisher Scientific). Cell lysates were separated on a 4-20% Mini-PROTEAN TGX Precast Protein Gels (#4561096; Biorad). Proteins were then transferred to PVDF membranes and membranes were incubated with antibodies to Sortilin-related receptor 1 (SORLA) at 1:1000 (# ab190684; abcam); Mouse monoclonal anti-Actin (clone A4) (#MAB1501; Millipore Sigma) at 1:2000; Mouse monoclonal anti-Phospho-Tau (AT8) (#MN1020; Thermo Fisher Scientific), Rabbit monoclonal anti-GAPDH (14C10) (#2118; Cell Signaling), Rabbit polyclonal anti-VPS35 ( #ab97545; Abcam) at 1:1000, Rabbit monoclonal anti-APP (#ab32136; Abcam) and Rabbit polyclonal anti-VPS26B (#NBP1-92575; Novus). For the PHF-Tau blots, blots were incubated overnight with PHF-1, Gift from Dr. Peter Davies, mouse 1:1000. After three washes with TBST, membranes were incubated with corresponding IRDye fluorescent secondary antibodies in intercept blocking buffer (Li-Cor) for 1h and scanned using an Odyssey Clx imaging system (Li-Cor).

**Immunocytochemistry**

Purified neurons were seeded at a density of 500,000 cells per well of a 24-well plate on glass coverslips coated with Matrigel. After 5 days in culture, cells were fixed in 4% paraformaldehyde (PFA, Alfa Aesar, Reston, VA) for 15 minutes. Cells were incubated in blocking buffer containing 2.5% bovine serum albumin and 0.1% Triton X-100 (Sigma Aldrich, St Louis, MO) for 30 minutes at room temperature then incubated in a primary antibody dilution in blocking buffer for 2 hours at room temperature. Cells were washed 3x with PBS + 0.1% Triton X-100 and incubated with a secondary antibody dilution in blocking buffer for 1 hour at room temperature. Cells were washed 3x in PBS and mounted on glass slides with ProLong Gold Antifade mountant (#P36930; Thermo Fisher Scientific, Waltham, MA). For details of antibodies used, see table S1.

**Confocal microscopy and Image processing**

Confocal z stacks were obtained using a Nikon A1R confocal microscope with x63 and x100 plan apochromat oil immersion objectives or a Yokogawa W1 spinning disk confocal microscope (Nikon) and a 100X plan apochromat oil immersion objective. Maximum intensity projections of confocal stacks were generated, and background was subtracted using the rolling ball algorithm. Endosome channels were enhanced using contrast limited adaptive histogram equalization algorithms (CLAHE) and masked using cell body stains. Size and intensity measurements were performed using Cell Profiler software(6). Individual puncta were identified using automated segmentation algorithms. Mean intensity of each puncta was measured and has been presented as a mean value over all puncta per field. Similarly, pixel area of each puncta was measured and has been presented as a mean area over all puncta per field. Finally, mean puncta area normalized by total cell area calculated from cell body stains is also presented

**Quantification and statistical analysis**

For early endosome size analysis, 3 clones of WT, 2 clones of E270K *SORL1*$^{var}$ , 1 clone of Y141C *SORL1*$^{var}$, 2 clones of G511R *SORL1*$^{var}$, 1 clone of *SORL1*$^{+/-}$ and 2 clones of *SORL1* KO were used. 10-15 images per clones were analyzed. For experiments measuring secreted Aβ, 3 clones of WT, 2 clones of E270K*SORL1*$^{var}$, 1 clones of Y141C *SORL1*$^{var}$, 2 clones of G511R *SORL1*$^{var}$, 1 clone of *SORL1*$^{+/-}$ and 1 clone of *SORL1* KO were used. 3 replicates per clone were analyzed for this experiment. For APP/EEA1 and VPS35/EEA1 colocalization experiments, 2 clones of WT, 2 clones of *SORL1* KO and 1 clone of *SORL1*$^{+/-}$ were used. 10 images per clone were analyzed for this experiment. For measurement of phosphorylated and total tau experiments, 2 clones of WT, 2 clones of E270K *SORL1*$^{var-}$, 1 clone of Y141C *SORL1*$^{var}$ , 2 clones of G511R *SORL1*$^{var}$, 1 clone of *SORL1*$^{+/-}$ and 2 clones of *SORL1* KO were used. 3 replicates per clone were analyzed for this experiment. For detection of tau using western blotting, 2 clones of WT and 2 clones of *SORL1* KO were used. 1-2 replicates per clone per condition were analyzed for this experiment. For early endosome size analysis with TPT-260 treatment experiments, 2 clones of WT, 2 clones of E270K *SORL1*$^{var}$, 1 clone of Y141C *SORL1*$^{var}$, 2 clones of G511R *SORL1*$^{var}$, 1 clone of *SORL1*$^{+/-}$ and 1 clone of *SORL1* KO were used. 15 images per clone per condition were analyzed for this experiment. For DQ-BSA assay, 1 clone of WT and 2 clones of *SORL1* KO were used and 10 images per clone per time point per condition was analyzed. For transferrin recycling assay, 2 WT clones, 1 SORL1$^{+/-}$, and 2 *SORL1* KO clones were used and 10 images per clone per time point per condition were analyzed.
